# Water Pollution and Administrative Division Adjustments: A Quasi-natural Experiment in Chaohu Lake of China

**DOI:** 10.1101/2021.08.23.457414

**Authors:** Jing Li, Di Liu, Mengyuan Cai

## Abstract

The adjustment of administrative divisions introduces a series of uncertain impacts on the social and economic development in the administrative region. Previous studies focused more on the economic effects of the adjustment of administrative divisions, while, in this paper, we also take environmental effects into consideration. The administrative division adjustment for Chaohu Lake is used as a quasi-natural experiment to explore the influence of the adjustment on pollution control. The synthetic control method is used in this study to access the effect of administrative division adjustment on the water quality indicators of Chaohu Lake and its internal mechanism. Some conclusions are as follows. First, after the administrative division adjustment, some water quality indicators, such as ammonia nitrogen, have indeed been alleviated; however, other major pollution indicators, such as chemical oxygen demand and dissolved oxygen, have deteriorated to varying degrees. Second, the results also reveal that improper development ideas, industrial excessive expansion, and the swing of economic growth and environmental goals are problems after the adjustment. Returning to the original intention of adjustment, rationalizing the Chaohu Lake management system and designing a sound and feasible accountability mechanism are fundamental measures to reduce pollution.

## 1. Introduction

The adjustment of administrative divisions is often used as an administrative tool to solve problems related to economic development or to stabilize conflicts between the central city and its neighboring cities (He et al.,2018). Fine-tuning of administrative areas can be used to solve regional problems and promote regional development. Regardless of military political factors or economic functions, a large number of natural experiments have been provided, for example, the merger of East and West Germany brought about by the fall of the Berlin Wall in 1990, the collapse of the Soviet Union in 1991, and the separation of North and South Korea caused by the Korean War in 1945. Moreover, in China, Hainan Province was established in 1988, and Chongqing Municipality was established in 1997. All these events have inspired discussion on impacts of administrative division adjustment. In Western countries, to promote regional coordinated development, the focus of administrative division has been on public services and government municipal cost, efficiency, and fairness (Boyne, 1995; Sancton, 1996). In contrast, the conflict in administrative division in China reflects the disaccord between the political system and economic reforms (Zhang and Wu, 2006). Since the reform and opening up in 1978, China’s administrative division has been constantly adjusted as part of the national strategy to promote rapid economic development (He et al.,2018) because China’s urban and regional development has been governed by territorial management in the transition from a planned economy to a socialist market economy (Ma, 2005). Consequently, there are obvious characteristics of the “administration area-delimited economy” in China.

The environment is a typical public good. Public goods and public services are nonexclusive and noncompetitive, which can easily lead to conflicts of interest between different local governments on environmental governance issues. This results in serious resource waste, environmental pollution, and damage. In recent years, increasingly serious problems, such as population explosion, excessive consumption of resources, environmental pollution, and ecological damage, have become major global issues (Chen, 2019), and environmental issues have received increasing attention. To cope with environmental governance issues in economic development, the 19th National Congress of the Communist Party of China proposed the “five in one” development concept and implemented the strictest environmental protection system. It is feasible to try to solve environmental problems in administrative regions by adjusting administrative divisions. However, due to its wide scope and complex content, research in this field is relatively limited, and the pertinence is relatively weak. Because of the numerous environmental factors and the difficulty of data acquisition, there are limited studies in this area, and no consistent conclusions have been reached. Some studies believe that the adjustment of administrative division will have a positive impact on environmental governance. They suggest that package planning of the regional environment through the adjustment of administrative divisions reduces the government’s buck-passing on environmental pollution control and clarifies the powers and responsibilities. Some scholars hold the opposite view, arguing that if authorities use frequent administrative division and administrative system adjustments to alleviate the contradiction between the economic foundation and the superstructure, they cannot establish a good regional governance model (Zhang and Wu, 2006) and cannot fundamentally eliminate economic problems of the administrative region. Even worse, it may instead be a legitimate reason for land expansion, leading to failure of environmental governance.

Chaohu Lake is the fifth largest freshwater lake in China and the largest lake in Anhui Province. Its water area covers approximately 825 square kilometers. Chaohu Lake is not only an important channel connecting the Jianghuai area to the north, but also an important agricultural product base. The Chaohu Lake basin covers a total area of 13350 square kilometers. Before the administrative division adjustment, it spanned 11 cities and counties, including Hefei, Chaohu, and Lu’an. Due to rapid social and economic development, the water quality of Chaohu Lake has gradually deteriorated. The pollution problem of Chaohu Lake has attracted the attention of the central and local governments. More than 80 billion yuan has been invested in treatment funds, but the pollution situation of Chaohu Lake has not been greatly improved. Before 2012, Chaohu Lake was jointly governed by Hefei and the prefecture-level Chaohu city. There was a lack of effective coordination between them, so pollution has not significantly alleviated, which has seriously restricted the regional economic development of the Chaohu Basin and affected the production and life of the local people. On August 22, 2011, with the approval of the State Council, the administrative division of Chaohu City was adjusted. The prefecture-level Chaohu city was officially rescinded into the county-level Chaohu city. One district and four counties under the jurisdiction of the original Chaohu city were placed under the jurisdiction of Hefei, Wuhu, and Ma’anshan. This made Chaohu Lake an inner lake of Hefei city, which facilitated the overall management of Chaohu Lake and provided development opportunities for Hefei. The effect of administrative division adjustment on the treatment of Chaohu Lake is considered in the quasi-natural experiment.

In this paper, the administrative division adjustment for Chaohu Lake is used as a quasi-natural experiment to explore the influence of the adjustment on pollution control. Using data from the Ministry of Environmental Protection, we choose the synthetic control method commonly used in project evaluation to identify the effect of administrative division adjustment on the water quality indicators of Chaohu Lake and discuss the internal mechanism. Our study contributes to the literature on environmental impacts of administrative division changes and is of great significance for understanding the environmental effects of administrative divisions and grasping the connotations of high-quality development.

## 2. Development of the Chaohu administrative division

The main driving force of urban development is market factors, but politics also has a subtle effect (Skinner, 1964). Since the reform and opening up in 1978, to meet the objectives of economic development, China’s administrative divisions have changed frequently. In contrast to the small-scale administrative division adjustments in foreign countries, those implemented in China often included transforming counties into cities and upgrading counties and cities in the early stage (Yu et al., 2018). In 2003, the Ministry of Civil Affairs of China raised the standard for transforming counties into urban districts, the large-scale administrative division adjustment gradually decreased, and the focus shifted to the adjustment of municipal districts and adjustments of districts and counties. Prefecture-level administrative division adjustments were mainly concentrated from 1978 to 2004, and targets ranged from the developed areas of the provinces to relatively developing areas. Therefore, phased adjustment was in line with regional economic development.

The administrative division adjustments of prefecture-level cities not only roughly constituted major urban agglomerations countrywide, but also created central cities and strengthened the functions of central cities. After 2004, adjustment of administrative divisions of prefecture-level cities was basically completed nationwide, and new changes were made with caution. Since 2000, the adjustments of administrative divisions have gradually shifted from city level to district level in China. Adjustment of large administrative districts at the prefecture level and above is relatively rare, while that of counties and municipal districts is more frequent (Figure 1). County-level administrative division adjustments can optimize a regional urban system, guide the direction of population migration and industrial agglomeration, and strengthen the economic and social ties among cities through the revocation, merging and upgrading administrative divisions (Yu et al., 2018). Internal adjustment of municipal districts can optimize and reconstruct the urban spatial structure, industrial layout, and public infrastructure configuration (Collin, 2002). Furthermore, improvements in administrative functions and management systems have also promoted the evolution of regional functions (Liu and Wong, 2018), thus forming the general pattern of administrative divisions in China.

**Figure 1.**
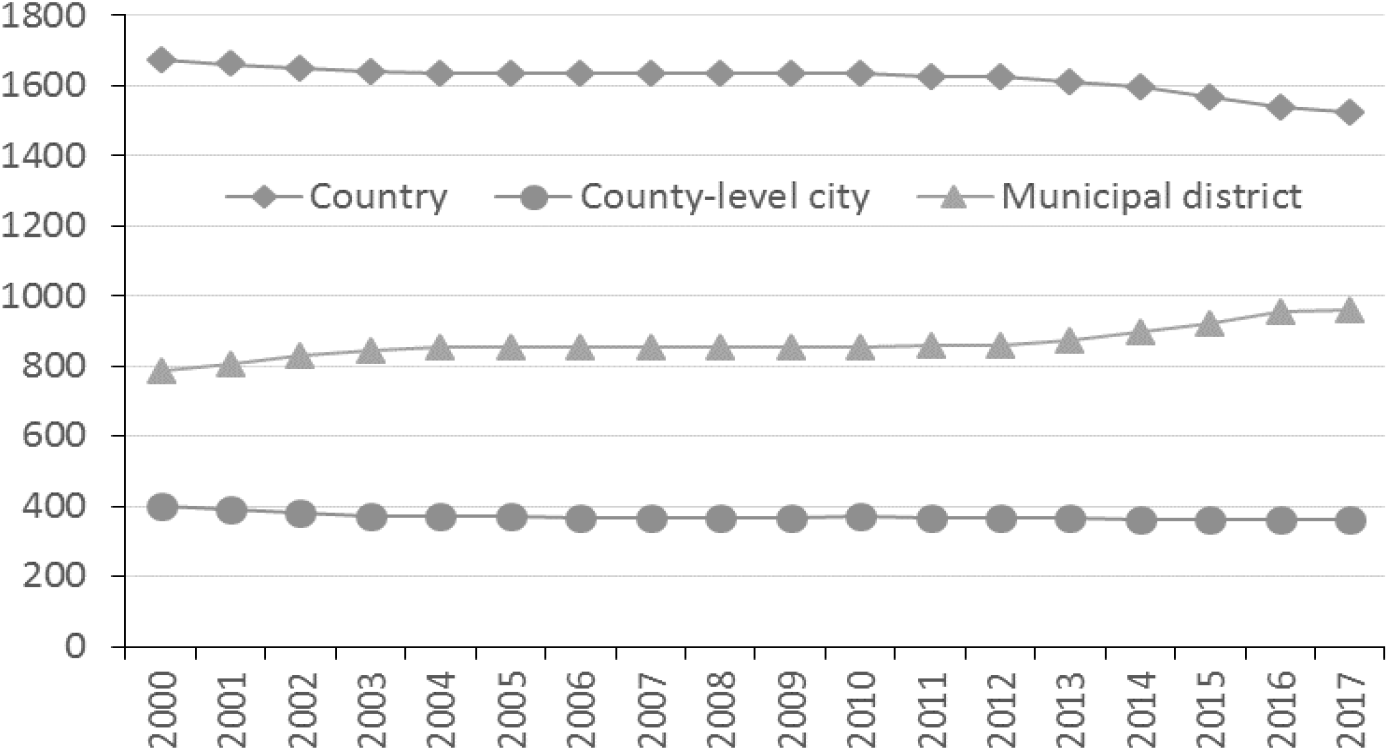
The numbers of administrative divisions of county level and above in China, 2000-2017

The original district-level Chaohu city consisted of one district and four counties. The city scale was small, and the level of economic development was relatively poor, while the difference among political districts was large, and problems related to river governance were severe. In particular, the water management system was not effective; hence, the pollution regulation of Chaohu Lake has not been fundamentally improved. To promote social and economic development and rationalize the management system of Chaohu Lake, in 2011, one district and four counties of the prefecture-level city were divided into three cities (see Figure 2). Lujiang was incorporated into Hefei, Hanshan and Hexian were incorporated into Ma’anshan, and Wuwei was incorporated into Wuhu.

**Figure 2.**
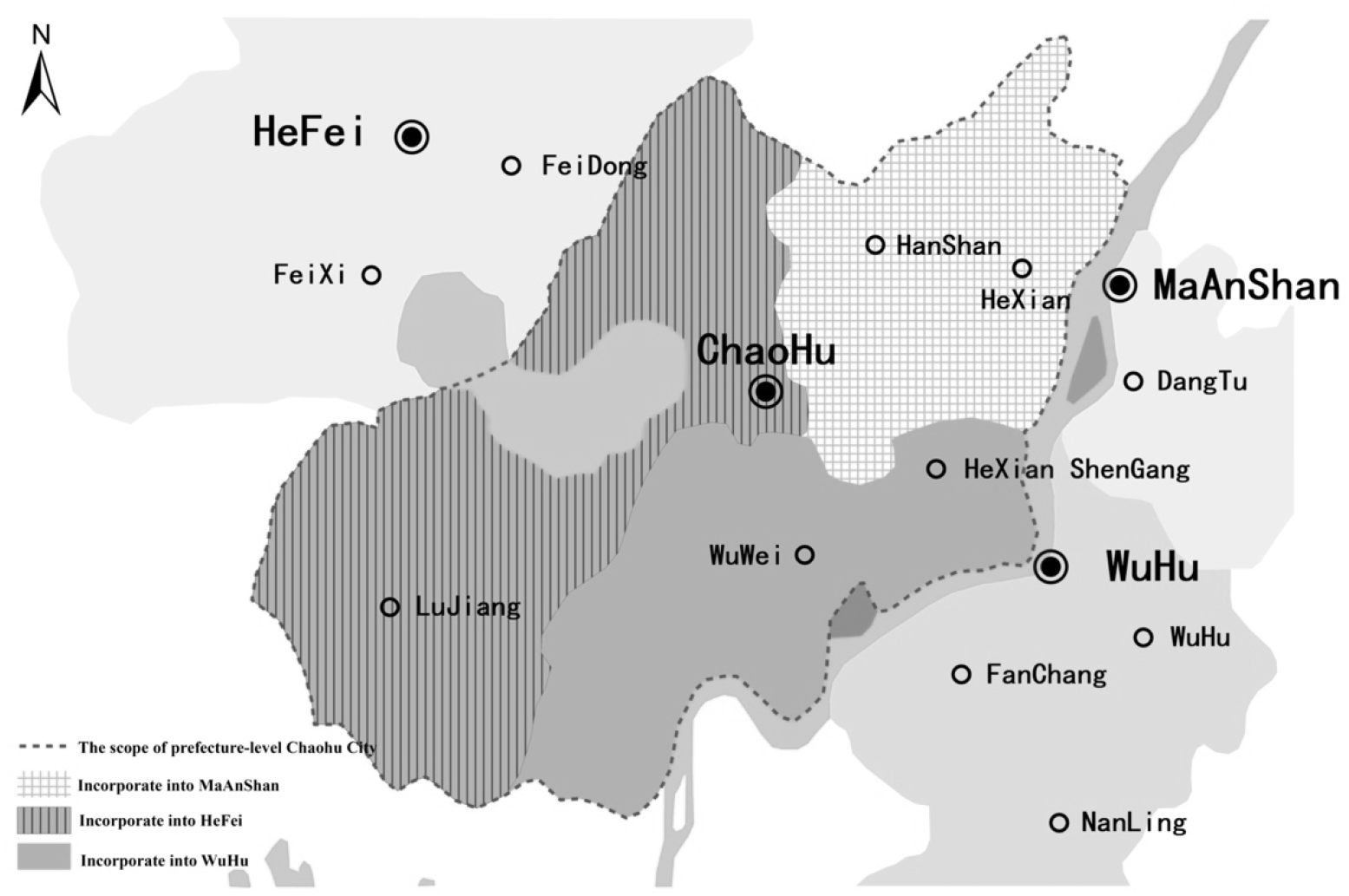
The scope of the prefecture-level municipality of Chaohu

The State Council’s approval of the separation of Chaohu Lake was based on economic factors and was originally intended to cope with contamination of Chaohu Lake. Prior to the adjustment, Chaohu Lake was governed by Hefei city and Chaohu city. The pollution control policies were difficult to unify, and the powers and responsibilities were unclear. As a result, externality of pollution was significant. However, after the adjustment, the Chaohu Administration Bureau was established to unify the management and planning of water conservancy, environmental protection, fishery administration, and other affairs, which was conducive to unifying the implementation of the ecological and environmental protection policies in Chaohu Lake Basin. In this way, problems caused by inconsistent enforcement of the environmental protection departments of the two places could be solved, and mutual shuffling events could be reduced. Hence, this change was an effective way to strengthen the comprehensive management of Chaohu Lake and enhance its capacity for sustainable development. Moreover, Hefei has stronger economic strength, which in theory is beneficial to the pollution control of Chaohu Lake. However, the strong interference in the ecosystem caused by urbanization, industrialization, and administrative division adjustment may cause the overall deterioration of ecological risks in the basin and pose a challenge to the comprehensive management of the Chaohu Lake water environment. The governance of large lakes is a worldwide intractable problem. Hefei must balance the relationship between rapid economic development and pollution control of Chaohu Lake and reasonably manage the industrial layout and ecological protection of Chaohu Lake Basin.

### 2.2 Environmental protection and water resources management system

China’s environmental protection system follows “territorial management and hierarchical responsibility”. The principle of territorial management requires relevant departments to manage environmental issues under their jurisdiction, which can strengthen the rights and responsibilities of local environmental protection departments in environmental supervision. However, the territoriality principle has disadvantages for rivers and lakes that cross administrative regions. Each department is responsible only for the effectiveness in its own region, which makes it difficult to implement comprehensive lake planning. Before the abolition of the prefecture-level municipality of Chaohu, Chaohu was jointly managed by Hefei city and Chaohu city and was divided into the east and west lakes for management. As a result, all departments considered the development and utilization planning only for their portion of the lake, and most administrative regulations and methods could not be effectively implemented. Disputes caused by regional joint governance are inevitable, and pollution control is difficult to achieve in such scenarios. After the abolition of the prefecture-level Chaohu city, Chaohu was under the independent jurisdiction of Hefei city, which avoided regional disputes caused by the joint governance of different administrative regions. Therefore, the abolition of the prefecture-level Chaohu city was conducive for Hefei to formulate independent overall planning and to clarify the responsibilities of government departments at all levels. In addition, compared with the prefecture-level Chaohu city, Hefei city has a developed economy, perfect urban functions, complete infrastructures, and a strong economic background. As a result, Hefei is able to analyze and study environmental carrying capacity of Chaohu Lake more scientifically, providing the possibility for better pollution control measures for Chaohu Lake.

For a long time, the assessment and promotion system of local governments has been based on economic growth, especially industrial growth and fiscal revenue. Use of the GDP growth rate as the main indicator in the promotion system has led to increased competition among local governments, which has reduced environmental standards in an effort to develop the economy. The promulgation of the newly revised Environmental Protection Law on April 24, 2014, proposed new requirements for local governments. It further clarified the governments’ supervision and management responsibilities for environmental protection and set penalties for officials who cannot perform their duties. Article 69 stipulates that leading cadres who report falsely or conceal the pollution situation will take the blame and be forced to resign. In the event of environmental violations that cause serious consequences, the leaders of the local government and the supervisors of the environmental protection department will bear the corresponding criminal responsibility. The new environmental protection law also specifies new requirements for enterprises. The amendment of the Environmental Protection Law includes very strict punishment measures: increased penalties for illegal sewage discharge companies that refuse to make corrections with continuous punishment on a daily basis without capping and fines on construction units that start construction without environmental protection approval. The implementation of the new environmental protection law places strict requirements on enterprises and local economic development, which is conducive to enterprises carrying out green production and to local governments developing a local economy without sacrificing the environment. These changes are also expected to be conducive to the improvement of the conditions of Chaohu Lake.

In addition, government has indeed made efforts to control Chaohu Lake. Table 1 shows that during these five-year planning periods, the government has invested considerable money to address the pollution problem of Chaohu Lake, and the number of completed projects has increased annually^①^. Among them, the 12^th^ Five-Year Plan (2011-2015) has invested four times as much as the 9^th^ Five-Year Plan (1996-2000). The Chaohu Basin Water Pollution Prevention Regulations have been continuously revised since they were adopted in 1998, and they were implemented in 2014 as a result of the twelfth meeting. Although the governance of Chaohu Lake is progressing slowly, targeted documents continue to set new standards for its governance. This process is conducive to the implementation of strategic deployment for the treatment of Chaohu Lake’s water pollution, improving the system, increasing the prevention and control of water pollution, and optimizing the social development environment of Chaohu Lake Basin. The administrative division adjustment has established the management system of Chaohu Lake basin management and administrative regional management, which is conducive to the meticulous design and overall promotion related to various aspects of the system, mechanism, and legal system to establish a veritable new system that is suitable for the sustainable utilization of water resources.

**Table 1.** Number of governance funds and projects invested in Chaohu Lake, 1996-2015.

| Period | Fund (Billion Yuan) | Number of Completed Projects | Pollution Situation |
| --- | --- | --- | --- |
| Ninth Five-Year Plan (1996-2000) | 2.58 | 3000+ | The progress of industrial wastewater treatment was slow, and the water pollution control did not reach the target. |
| Tenth Five-Year Plan (2001-2005) | 3.03 | 26 | Pollutant emissions did not meet planned targets. |
| Eleventh Five-Year Plan (2006-2010) | 7.07 | 56 | Due to the need of the Chaohu Lake Basin to undertake the demonstration zone of industrial transfer in the Wanjiang Urban Belt, the amount of industrial wastewater and pollutants also increased by more than 30%, and the pressure on environment risk prevention increased. |
| Twelfth Five-Year Plan (2011-2015) | 10.9 | 117 | Water quality in Chaohu Lake has been improved, but overall eutrophication status has not fundamentally changed. |

Note: The data are extracted from *The 10<sup>th</sup>-12<sup>th</sup> Five-Year Plan for Water Pollution Prevention and Control in Chaohu Lake Basin*, and *The Overall Plan for Water Environment Treatment of Chaohu Lake Anhui Province*.

### 2.3 The conflict between local development and environmental protection goals

The assessment and promotion system of local governments has always been based on GDP, an economic performance indicator, which has led local governments to focus excessively on the GDP growth rate and to lower the threshold of resource supply and environmental standards. To enhance their own competitiveness, local governments pay attention to economic growth and urbanization development but mostly ignore the carrying capacity of the natural ecological environment. Local governments attempt to avoid the regulation of environmental system in environmental management and seek approval of economic projects, investment, and operation. The rapid expansion of the urban scale of Hefei has caused contradictions between resources and urban development. The new plan will inevitably cause friction between the economic and environmental goals of the original district. Figure 2 shows that the area under the jurisdiction of Great Hefei has suddenly expanded by 4,379 square kilometers, forming a new development pattern of the “surrounding Chaohu Lake and connecting the Yangtze River to the south”. From the perspective of administrative area, Hefei has already become a regional mega-city; however, in terms of environmental governance, Hefei has not given full play to the advantages of Chaohu Lake governance caused by the adjustment of administrative divisions.

After the abolition of the prefecture-level municipality of Chaohu, Hefei gained a geographical space for development and paid more attention to economic development: Chaohu Lake governance was focused on assisting industrial development. The rapid development of Hefei in recent years has led to a sharp increase in the scale of river basin construction and rapid population growth, and the continuous accumulation of pollution in production and life has far exceeded the carrying capacity of Chaohu itself. Although the causes of pollution are obvious, it has not been taken into consideration in the process of development and construction around Chaohu Lake in recent years. Environmental supervision^①^ has noted that in the context of the rapid economic and social development of the Chaohu Basin, the water pollution in Chaohu Lake has shown a trend of improvement, but the “Regulations on the Prevention and Control of Water Pollution in the Chaohu Lake Basin” has not been successfully implemented. The division of the first, second, and third protection areas of Chaohu Lake is one of the core provisions of the most stringent regulation in history, but this work remains in the expression of basic principles stage when it should have become the safety bottom line of Chaohu protection and has been put on the shelf. Whether the regulations can be enforced is essentially a question of who gives way when there is a conflict between environmental protection and development: pollution control frequently gives way to economic development in Chaohu.

Table 2 shows that the eutrophication status of the western half of Chaohu Lake was generally worse than the eastern half. From the perspective of comprehensive eutrophication state index, the data did not change substantially, and as a whole, they did not show a positive trend. Although Hefei has invested considerable amounts of money, the existing Chaohu Lake governance focuses on road construction, river regulation, and comprehensive ecological measures. The eutrophication status of Chaohu Lake has not fundamentally improved. In recent years, with the high incidence of water blooms, the cyanobacteria data reported by the environmental supervision group highlight the urgency of pollution control. In 2015, the largest bloom area was 321.8 square kilometers, accounting for 42.2% of the whole lake, the highest in nearly eight years, while it was 237.6 square kilometers in 2016, accounting for 31.2% of the entire lake.

**Table 2.** Eutrophication status of Chaohu Lake from 2009 to 2016

| Year | Comprehensive Eutrophication State Index |  |  | Eutrophication Status |  |  |
| --- | --- | --- | --- | --- | --- | --- |
|  | Eastern-half | Western-half | Whole | Eastern-half | Western-half | Whole |
| 2009 | 52.3 | 64.8 | 60.1 | LE | SE | ME |
| 2010 | 51.0 | 64.2 | 59.0 | LE | ME | LE |
| 2011 | 52.3 | 63.7 | 59.5 | LE | SE | ME |
| 2012 | 53.5 | 60.9 | 57.4 | LE | ME | LE |
| 2013 | 52.2 | 60.0 | 55.8 | LE | ME | LE |
| 2014 | 50.5 | 63.1 | 57.1 | LE | SE | ME |
| 2015 | 50.6 | 62.8 | 57.2 | LE | ME | LE |
| 2016 | 50.1 | 62.5 | 56.8 | LE | ME | LE |

Note: LC: Light Eutrophication; ME: Medium Eutrophication SE: Serious Eutrophication.
The data are extracted from the "China Environment Yearbook 2010-2017" and the National Surface Water Monitoring Station of the Ministry of Environmental Protection.

## 3. Literature review

In general, several significant impacts are considered in assessment of administrative division adjustment. The more common effects concern economics, which are the focus in many previous studies, while a small number of studies evaluate environmental effects of administrative division adjustment. A brief literature review of the two major effects is included in the following discussion.

### 3.1 A review of literature on economic effects of administrative division adjustments

Some studies have examined the impacts of administrative division adjustments on local economic development in terms of social and political systems and economic development stages. For instance, Wagenaar et al. (2004) found significant correlations between administrative division adjustments and local economic growth in the US. Ideally, administrative division adjustments could lead to substantial changes in local economic development. In the US, administrative division adjustments are often implemented to control the spread of diseases and protect local ecology, but rarely for economic development considerations. A wave of merging cities to form metropolises in Canada during the 1990s primarily aimed to reduce administrative costs and improve the efficiency of public services (Downey, 1998). The mergers overcame administrative boundary barriers to integrated local development, especially declines in core cities, and generated remarkable positive effects. Redding et al. (2008) considered the German division and reunification as a natural experiment and studied whether strong correlations exist between the local economic development of border cities and administrative division adjustments. Similarly, Abadie et al. (2015) assessed the impacts of reunification on the per capita gross domestic product (GDP) in West Germany. They found that the impacts of administrative division adjustments due to German reunification emerged in 1992 and that East Germany was a burden on West Germany’s economic development.

Due to the special features of China’s economic system, administrative division adjustments are often implemented for economic considerations. Administrative division adjustments are a major guiding factor of urbanization in China and have far-reaching economic and social impacts on local development (Yang, 2017).

Vast literature on this topic exists, and extensive studies on administrative division adjustments in China generally fall into two groups. The first group is based on case studies of administrative division adjustments in a specific region (province/municipality) and compares the local economic volume and economic growth before and after the adjustments. For instance, Chen (2006) studied administrative division adjustments in Sichuan province between 1993 and 1998. The results revealed that the administrative division adjustments indeed boosted Sichuan’s economic growth. Zhang and Wu (2006) describe conflicts in the Yangtze Delta. Due to administrative division adjustments and large-scale merging, local governments’ political interventions in economic activities have been intensifying since 2000. Administrative division adjustments, in fact, reflect disharmony between political systems and economic development. The second group analyzes the causes, patterns, impacts, and reform approaches of administrative division adjustments in China. The historical establishment of administrative divisions is based on national unity and political stability, and modern administrative division adjustments are implemented mainly to address the evolving needs of rapid economic and social development and urbanization (Zhou, 2008; Zheng and Kahn,2013; Wang and Liu, 2015). Yu et al. (2018) provided a comprehensive summary of administrative division adjustments at the province and prefecture levels in China. They concluded that administrative division adjustments have been a major factor influencing the progress of urbanization progress and a substantial driving force for the evolution of the urban spatial structure in China.

In general, the existing literature on how administrative division adjustments affect local economic development can be classified into two categories. The first category focuses on the overall influences of administrative division adjustments on local economic development. Due to variations in development levels of different regions and different administrative division adjustment approaches, adjustments are implemented in different ways. Moreover, effects of adjustments on economic development are not always positive and vary from region to region. The other category focuses on analyzing how administrative division adjustments affect a specific aspect of the local economy, such as industrial upgrades, urbanization, and interregional inequality. More studies fall into this category, with more specific analysis indicators, and the results vary greatly from each study.

### 3.2 Review of the environmental effects of administrative division adjustments

Chinese and overseas scholars have conducted studies on the environmental governance impacts of administrative division adjustments. Most existing studies have considered economic impacts as the primary study subject and covered changes in environmental aspects only in passing, or they includedfew qualitative assessments on the environmental impacts of the adjustments: empirical assessments are very rare. Studies from abroad focus on the transboundary transfer of pollution. The environmental standards and policies of different administrative divisions can cause a spatial reallocation of pollution, as lax environmental regulations can be effective in attracting current capital, and emissions reduction can deter international investments (Becker and Henderson, 2000; Wolfgang and Levinson, 2002; List et al., 2003). Consequently, polluting enterprises move their operations from one administrative division to another to avoid requirements for pollution control. Duvivier and Hang (2013) noted that regardless of whether transboundary shifting of pollution exists in Hebei province, the regions at provincial borders are more attractive than their counterparts in the hinterland among polluting enterprises. Similarly, Cai et al. (2016) discovered that polluting enterprises often prefer moving to administrative borders. Therefore, pollution shifting among different administrative boundaries is a major barrier in eliminating environmental pollution. Administrative division adjustments are important instruments for addressing environmental issues. Due to different social and political systems and economic development stages, there are fewer studies on administrative division adjustments overseas, and the adjustments are implemented mainly to protect ecological systems and prevent the spread of disease.

Water is a core resource in sustainable development and is critical for socioeconomic development, ecological systems, and human survival (Yao et al., 2016). Water resources involve many complex factors and correlations, including some interrelated natural, social, and economic elements. In the process of water resource utilization, complicated interrelations subject water resources to the influences of many uncertainties from multiple aspects (Chen et al.,2019). Chinese local governments can exert administrative power only within their administrative divisions; the transboundary feature of pollution makes it impossible to hold a specific local government accountable (Chen et al.,2018). The increasing competition and conflicts among different regions and sectors for water use enhance the significance of rational water allocation and utilization (Jaramillo and Nazemi,2018; Lipscomb and Mobarak, 2017). Consequently, most environmental regulations cover multiple dimensions and are difficult to quantify (Shadbegian and Wolverton, 2010). Since 1978, China has implemented a pollution control and prevention system based on a division of power. The central government centralizes the power of the overall environmental target setting, while local governments are responsible for formulating and implementing specific environmental regulations (Van Rooij and Lo, 2010). Until 2005, local officials lacked strong motives to realize the environmental targets set by the central government, and the career development of local officials was entirely dependent on local economic growth rates during their tenures (Zhou, 2008). Therefore, environmental degradation was common and persistent (Vennemo,2009).

To ensure that local governments comply with the environmental targets set in the 11th Five-Year Plan (2006-2010) released in 2005, the central government revised the performance assessment criteria for local officials, making performance in achieving pollution control targets a key indicator in determining the career development of local officials (Chen et al., 2018). Local officials who fail to achieve pollution control targets can be fired (State Council, 2007). The new system gives local officials a strong incentive to accomplish national environmental targets (Kahn et al., 2015). Chinese scholars have not conducted many studies on the environmental governance effects of administrative division adjustments. Instead, they tend to conduct simple analyses of the environmental governance impacts while studying the overall impacts of administrative division adjustments or employ administrative division adjustments as one of the factors influencing a specific region’s ecological and environmental governance changes during a specific period.

## 4. Methods

To investigate how the abolition of the prefecture-level municipality of Chaohu could affect water quality in Chaohu Lake in the Hefei region, analysis would typically have to be performed under a counterfactual framework. Factors determining environmental governance vary in different cities, and factors affecting water quality of rivers and lakes are complicated even without division adjustments. Therefore, the hypothesis of convergence among individual samples under the double-difference method cannot be satisfied. The matching method does not work neither, as it is impossible to find a region that is similar to the Hefei region in all aspects that did not undergo administrative division adjustments. Moreover, in terms of geographic, historical, regional, and other factors, different lakes have different levels of water pollution, and it is impossible to find a lake that is similar to Chaohu Lake. Due to these considerations, this paper adopts the synthetic control method proposed by Abadie et al. (2003). To assess the effects of an event, Rubin’s counterfactual framework is established first to represent how the area would be in the absence of a specific event (counterfactual); then, the data under the counterfactual scenario are compared with the actual data after the event occurred. The differences between the two scenarios are known as treatment effects. Constructing data for a control group that is similar to the region is an important exercise. To create a control group using the synthetic control method, the comparison is not made with one or multiple similar regions; instead, a reasonable control group is created based on a weighted average of all similar regions. The property variables for the control group are almost identical to those for the treatment group before the event and represent the status of the region in the absence of the event. The synthetic control method has three major advantages in the establishment of a counterfactual framework. First, when designing the control group by calculating the weighted average of similar regions, the weights are decided by the data, not by subjective choices; each weight represents the region’s contribution to the control group. Second, the sum of the weights of all samples is equal to one, and such weights avoid excessive extrapolation. Third, the synthetic control method requires that each component region of the control group is similar to the treatment region. For this reason, highly different regions should be excluded in the control group (Temple, 1999).

The study considers the abolition of the prefecture-level municipality of Chaohu as a treatment event and assesses its effects on the water quality of Chaohu Lake in the Hefei region. Assume time *t* ∈ [1,*T*], and let the year of administrative division adjustment be *T*_0_. 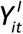 and 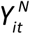 represent that region *i* was affected and was not affected by administrative division adjustment at time *t*. Then, 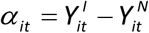 can represent the effects of the administrative division adjustment. In the case of the Hefei region, when *T*_0_ < *t* ≤ *T*, i.e., after abolition of the prefecture-level municipality of Chaohu, 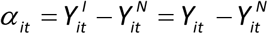, where *Y_it_* represents the actual situation in Hefei, while 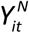 is a counterfactual value, i.e., the situation in Hefei would have been in the absence of the administrative division adjustment.

To observe the administrative division adjustment of the municipality of Chaohu, a total of J+1 samples is distributed into the treatment group (Hefei region) and control group. The factor model as proposed by Abadie et al. (2010) is used to synthesize 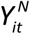, and the weights for the control group are defined as *w* = (*w*_2_, *w*_3_,…,*w_J+1_*)′, where *w_J_* is the weight of synthetic control city J. For any given *w_J_*, the synthetic control city can be calculated as follows:

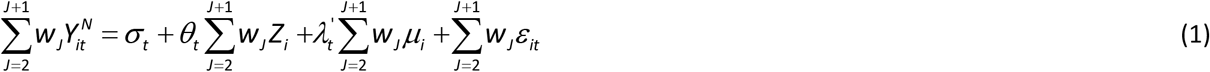

where σ_*t*_ represents the time-independent effect of the abolition of the prefecture-level municipality of Chaohu, *Z_i_* is the observed vector, *θ_t_* is the unknown coefficient, 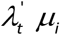 are the unobservable interactive fixed effects, 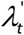 represents common unobservable factors, *μ_i_* is the unobservable regional fixed effects, and *ε_it_* is a random error term.

When the intervention period *T*_0_ is close to infinite, the optimal *w*\* should exist, and the synthetic control estimator is asymptotically unbiased. However, such conditions do not occur in reality. Usually, unobservable characteristics of the synthetic control group are made similar to those of the treatment group. Abadie et al. (2010) demonstrated that when *t* ≤ *T*_0_, if *w* that enables both 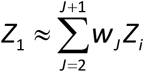 and 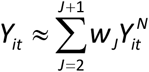 can be found, then it could also lead to 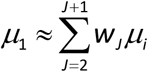; therefore, the evolution pathway of the effect of the synthetic Hefei region is similar to the actual growth pathway of Hefei before the abolition of the municipality of Chaohu. After calculating the weight matrix using data before the administrative division adjustment, 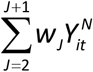 can be used as an unbiased estimator of 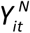. Hence, an unbiased estimate of *α_it_* can be obtained as follows:

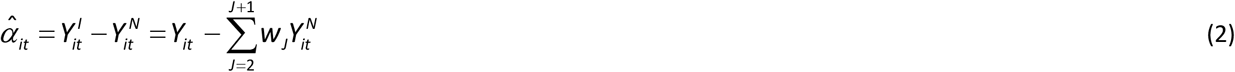

where 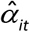 represents the environmental and economic effects of the abolition of the prefecture-level municipality of Chaohu, *α_it_* > 0 indicates that the effects of the administrative division adjustment are positive, and *α_it_* < 0 indicates that the effects are negative.

## 5. Empirical results and analysis

The results of the synthetic control of the environmental effects of Chaohu Lake are discussed in this section, followed by several robustness tests.

### 5.1 Results of the synthetic control of the environmental effects of Chaohu Lake

In the synthetic control method estimation, Hefei is used as the experimental group, and the remaining 69 samples are selected as the synthetic Hefei control group. The weights of 69 sample points are mostly non-zero, which ensures a comprehensive adoption of samples. Moreover, samples that are more similar to Hefei are assigned with larger weights. The chemical oxygen demand (COD), ammonia nitrogen (NH_3_N), and dissolved oxygen (DO) indicators are used to construct Hefei and synthetic Hefei’s indicator paths and their predictors. COD reflects the degree of organic pollution in water, and a higher COD is regarded to a more serious pollution. NH_3_N is derived from synthetic chemical fertilizers and is usually related to agricultural emissions. Additionally, DO reflects water pollution, especially organic pollution. The variables of the study are mainly divided into two categories: environmental and economic data, as shown in Table 3. The environmental data is extracted from the national surface water monitoring station of the Ministry of Environmental Protection. The annual average data is reorganized and matched with the economic and social data of the city where the monitoring point is located. Because the SCM requires completely balanced panel data, some missing samples are deleted, and finally a panel data set of 70 samples covers four types of water quality data and six types of socio-economic indicators. The economic data are mainly compiled from the China City Statistical Yearbook and Provincial Statistical Yearbook. We ensure that the synthetic control group is sufficiently close to the characteristics of synthetic Hefei, Tibet, and some prefecture level city data are excluded from the study. Table 3 shows the descriptive statistics of the environmental and economic variables.

**Table 3.**
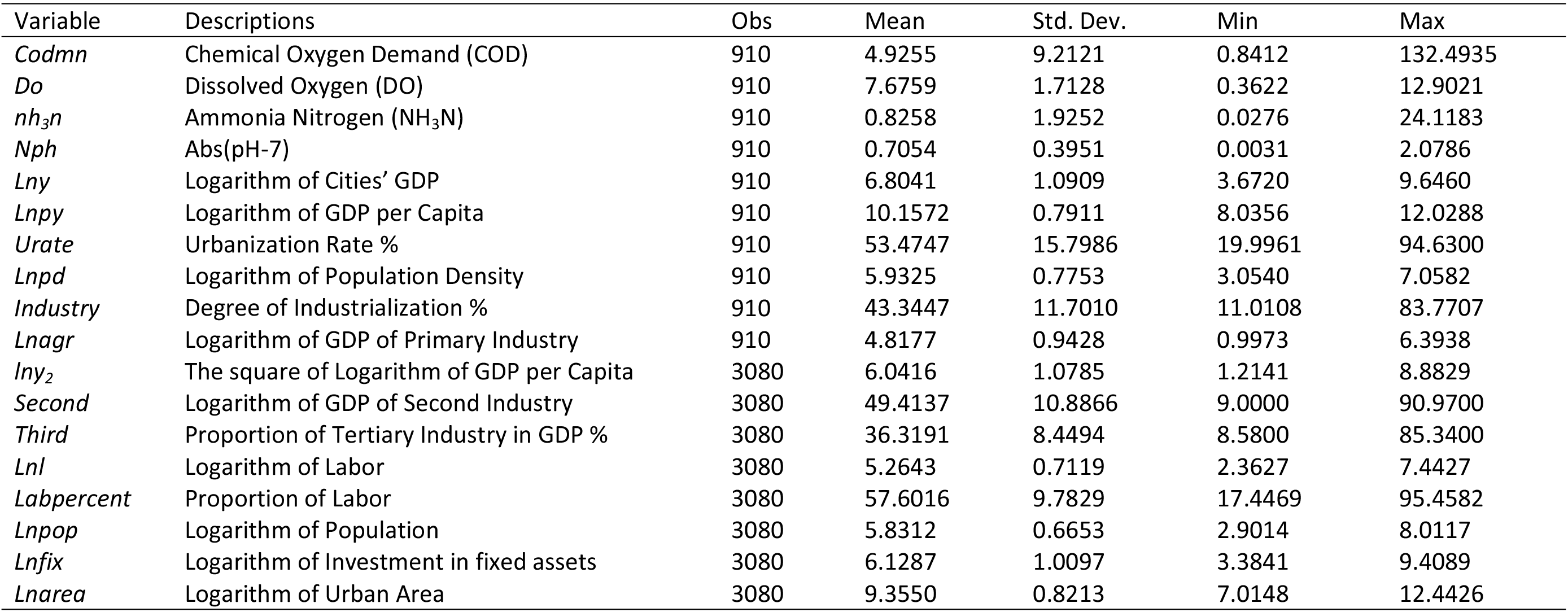
Descriptive Statistics

Figure 3 shows that prior to the abolition of the prefecture-level municipality of Chaohu in 2011, the CODs of synthetic Hefei and Hefei coincide to a high degree, which indicates that synthetic Hefei is a good counterfactual. After the abolition of prefecture-level municipality of Chaohu, the COD indices of Hefei and synthetic Hefei display different trends. The COD index of synthetic Hefei continues a downward trend, while that of Hefei shows an upward trend. This difference is mainly due to a series of rapid urban construction and industrial development projects implemented by the newly formed Binhu New District to Chaohu city and Lujiang county, leading to a large amount of organic pollution in Chaohu Lake, shown as a sharp rise in the COD index.

**Figure 3.**
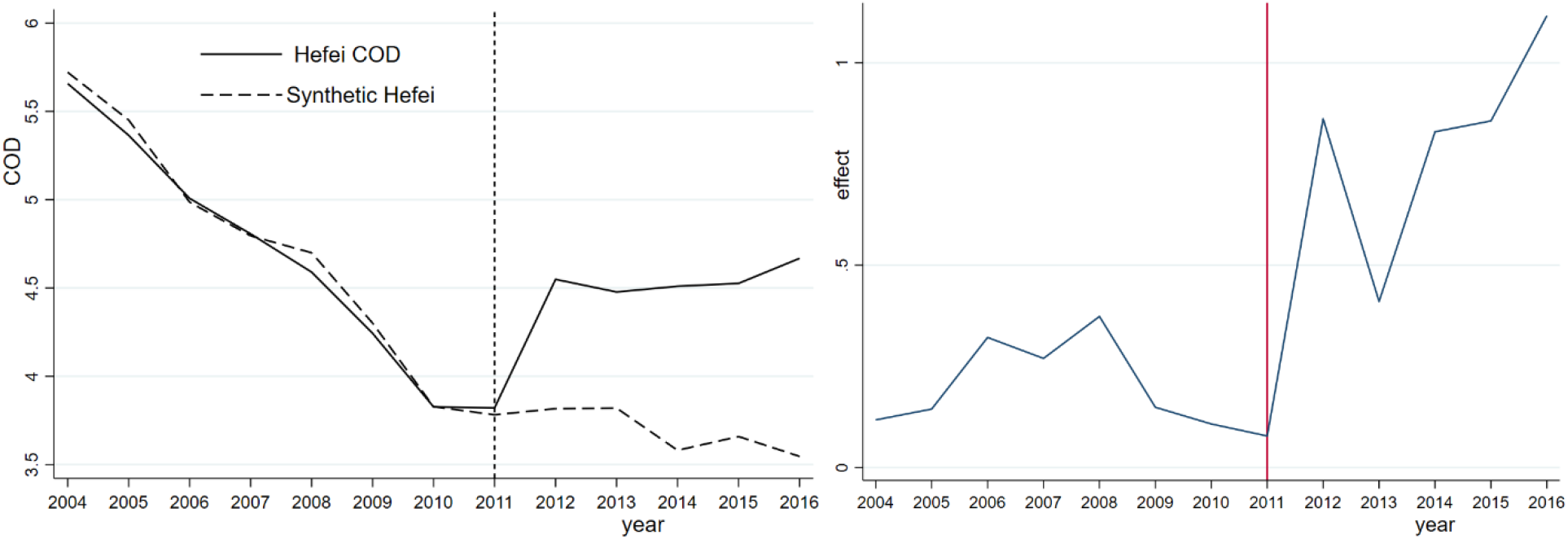
Hefei synthetic COD and its effect

The annual COD difference between Hefei and synthetic Hefei is calculated to assess the effect of administrative division adjustment on COD in Hefei. Before the adjustment, the COD gap between Hefei and synthetic Hefei is very small, but it increases after the abolition of the prefecture-level municipality of Chaohu in 2011. On the one hand, the COD index of synthetic Hefei matches that of Hefei, on the other hand, the adjustment of Chaohu city causes a higher COD level in Hefei. Table 4 compares the predictions of the COD index for Hefei and synthetic Hefei before the adjustment of the prefecture-level Chaohu urban area in 2011. From Table 4, we observe that GDP, GDP per capita, GDP of the secondary industry, urbanization rate, and population density are similar between the treatment group and the synthetic group.

**Table 4.**
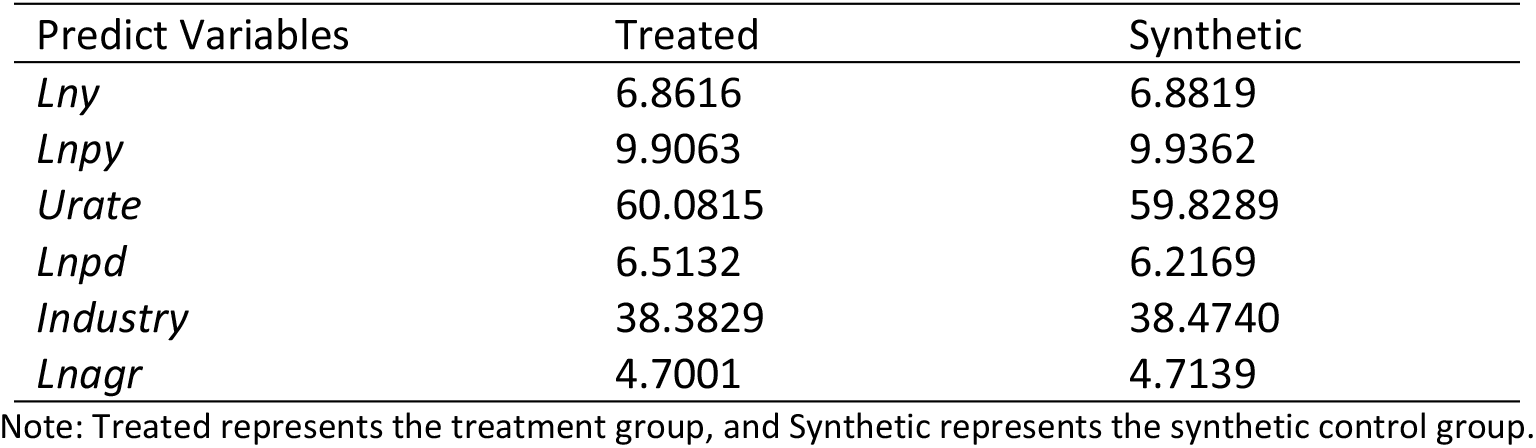
Treatment and synthetic control group of predictive variables for COD

In contrast, the synthesized NH_3_N shows a steady downward trend, but the actual NH_3_N index declines sharply after adjustment of the division, implying a significant improvement in water quality. As previously noted, NH_3_N originates mainly from agricultural emissions rather than industrial emissions and is the result of rapid urban development in Hefei, which reduces the amount of land surrounding Chaohu Lake used for agricultural purposes. As a result, the use of chemical fertilizers and pesticides has declined significantly, directly reducing the pollutants. From Figure 4, we observe a large gap between the NH_3_N indices of Hefei and synthetic Hefei during the 2011-2013 period. However, the gap gradually narrows, and the continuous improvement in NH_3_N in Hefei is not obvious after 2013. As reported in Table 5, the fitting error of each variable between the treatment group and synthetic control group is relatively small, that is, less than 0.1. In other words, synthetic Hefei is a good counterfactual for Hefei in terms of the ammonia nitrogen index.

**Figure 4.**
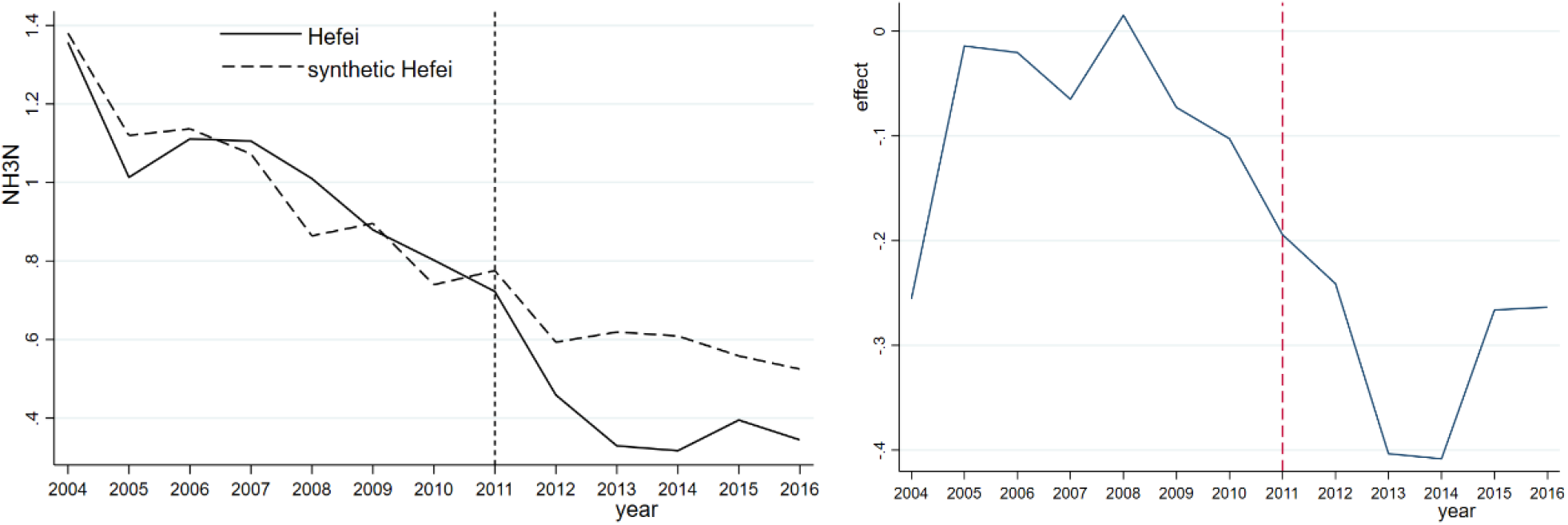
NH_3_N and its effect for Hefei and synthetic Hefei

**Table 5.** Treatment and synthetic control group of predictive variables for NH_3_N

| Predict Variables | Treated | Synthetic |
| --- | --- | --- |
| *Lny* | 6.8616 | 6.8502 |
| *Lnp<sub>y</sub>* | 9.9063 | 9.8948 |
| *Urate* | 60.0815 | 60.0211 |
| *Lnp<sub>d</sub>* | 6.5132 | 6.5062 |
| *Industry* | 38.3829 | 38.3609 |
| *Lnagr* | 4.7001 | 4.6916 |

Note: Treated represents the treatment group, and Synthetic represents the synthetic control group.

In terms of the DO indices, as illustrated in Figure 5, similar levels of DO are observed for Hefei and synthetic Hefei prior to 2011. Although the synthetic DO index increases annually from 2011 to 2016, there is no obvious improvement in the DO index in Hefei. Similar to COD, the DO index is not improved due to the vigorous development of the shores of Chaohu Lake and the construction of a large number of buildings without corresponding environmental protection measures. Unsurprisingly, the DO index of Hefei is always lower than that of synthetic Hefei, especially one year after the adjustment. Table 6 compares the predicted DO levels of Hefei and synthetic Hefei before abolition of the prefecture-level municipality of Chaohu in 2011. The regression results for each variable indicate that the difference between the synthetic Hefei and Hefei city is very small, indicating that a comparison of the fitting effect is appropriate.

**Figure 5.**
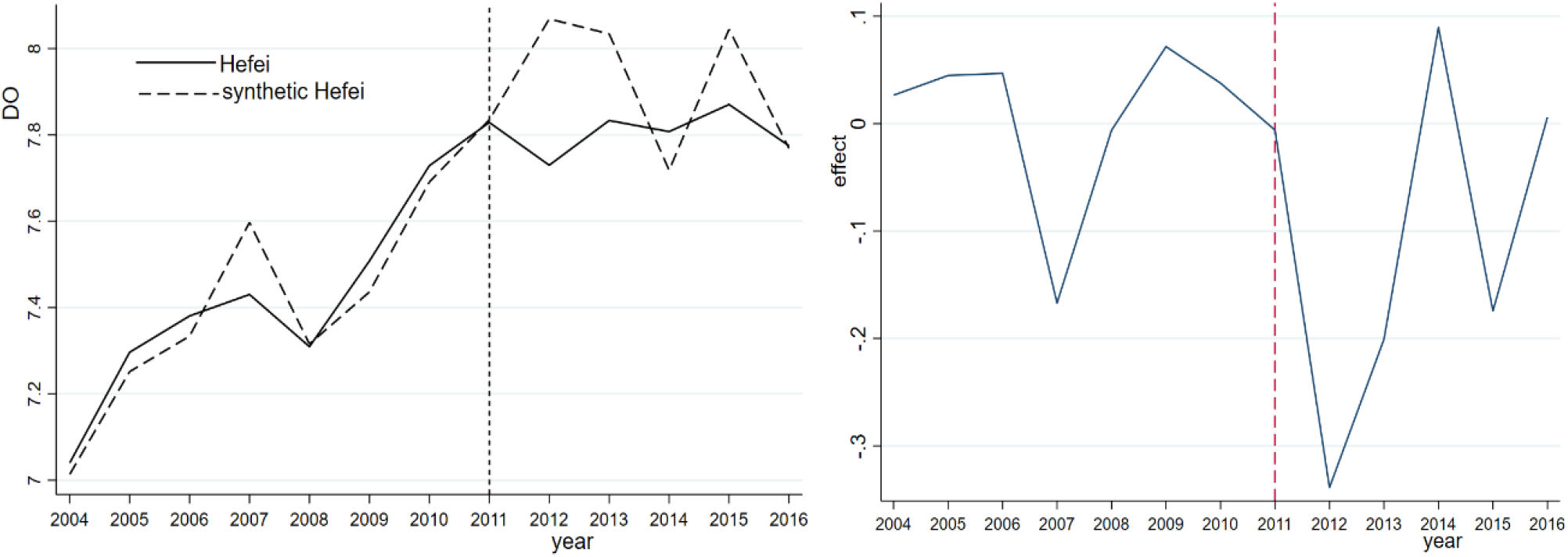
DO and its effect for Hefei and synthetic Hefei

**Table 6.** Treatment and synthetic control group of predictive variables for DO

| Predict Variables | Treated | Synthetic |
| --- | --- | --- |
| *Lny* | 6.8616 | 6.8491 |
| *Lnp<sub>y</sub>* | 9.9063 | 9.8864 |
| *Urate* | 60.0815 | 60.0184 |
| *Lnp<sub>d</sub>* | 6.5132 | 6.4997 |
| *Industry* | 38.3829 | 38.2643 |
| *Lnagr* | 4.7001 | 4.6896 |

Note: Treated represents the treatment group, and Synthetic represents the synthetic control group.

### 5.2 Robustness check

To validate the results, several robustness checks are conducted in the study. First, the DID method is the most intuitive method in policy evaluation and can be used to compare changes in the water quality of Chaohu Lake after the administrative division adjustment and changes in the water quality at other monitoring points. Table 7 shows the regression results using the DID method.

**Table 7.** DID method regression results

| Variables | COD | NH<sub>3</sub>N | DO |
| --- | --- | --- | --- |
| *chaoxt* | -7.878**<br>(3.747) | -3.347***<br>(0.683) | 1.015<br>(0.980) |
| *Ln<sub>y</sub>* | 11.04***<br>(3.801) | 3.341***<br>(0.645) | -3.055***<br>(0.780) |
| *Ln<sub>py</sub>* | -3.352<br>(2.790) | -1.501***<br>(0.486) | 3.185***<br>(0.697) |
| *Urate* | -0.267***<br>(0.0892) | -0.0224**<br>(0.0108) | -0.0226<br>(0.0180) |
| *Ln<sub>pd</sub>* | 7.040*<br>(4.250) | -0.185<br>(0.855) | -0.998<br>(1.241) |
| *Industry* | 0.243**<br>(0.115) | 0.0163<br>(0.0156) | -0.0100<br>(0.0111) |
| *Ln<sub>agr</sub>* | -4.131**<br>(1.736) | 0.232<br>(0.314) | 0.388<br>(0.369) |
| Year Fixed Effect | Yes | Yes | Yes |
| City Fixed Effect | Yes | Yes | Yes |
| Monitoring Point Fixed Effect | Yes | Yes | Yes |
| River System Fixed Effect | Yes | Yes | Yes |
| Constant | -59.38**<br>(29.32) | -8.301<br>(6.581) | 5.835<br>(9.671) |
| Obs | 910 | 910 | 910 |
| R<sup>2</sup> | 0.700 | 0.808 | 0.703 |

Note: The robust standard error (in parentheses); \*\*\* means $p < 0.01$ , \*\* means $p < 0.05$ , \* means $p < 0.1$ .
*chaoxt* is a policy variable where *chao* is a dummy variable that is equal to 1 if the Chaohu Lake observation point is observed, and *t* is a dummy variable that is equal to 1 after the administrative division adjustment.

1. Prefecture-level cities are used in the study. The municipalities in Beijing, Shanghai, and Tianjin are significantly different from those in other provinces, such as Hefei, in terms of economic characteristics and other aspects. Therefore, the data for the three municipalities in Beijing, Shanghai, and Tianjin are excluded in constructing a control group of synthetic Hefei for these three indicators. The regression results, as shown in Table 7, confirm that the new synthetic and synthetic Hefei are roughly similar. For instance, the administrative division adjustment improves the NH_3_N index but not the COD and DO indices (Figures 6–8).
2. From Table 7, we observe that changes in COD and NH_3_N are significant at the 5% and 1% levels, respectively, while the change in DO index is not significantly different from zero. However, the DID method is not appropriate for evaluating administrative division adjustment policies. This is because DID has three major problems when addressing the possible water quality impacts of administrative division adjustment (Bertrand et al., 2004). (a) The selection of the reference group is subjective and arbitrary. (b) The DID requires that the experimental group and control group have the same trend before implementation of the policy, that is, the common trend hypothesis. However, the experimental group has only one observation, i.e., Hefei, and it is difficult to satisfy this assumption. (c) The problem is endogenous. In many cases, this endogeneity cannot be ruled out, and direct estimation using DID will produce biased results. In contrast, our synthetic control method is more robust to common factor shocks and nonlinearities compared with DID method.
3. Following Abadie and Gardeazabal (2003) and Abadie et al. (2010), a placebo test is conducted for robustness purposes. Ideally, the study will identify a place without division adjustment and compare the water quality index between synthetic Hefei and Hefei. Since weights are important for synthetic variables, the two observation points with the largest and smallest weights in the control group (i.e., Xinchengqiao and Haibowan) are tested with a placebo to observe changes in their water quality indicators. Figure 9 shows the observations with the largest and smallest weights in the synthetic control group. If the gap is observed, then the synthetic control method has failed to provide sufficient evidence that the administrative division adjustment affects the water quality of Chaohu Lake. From Figure 9, there is no difference between the actual and synthetic water quality indices, indicating that there is no obvious policy impact on the synthetic effect of the Xinchengqiao and Haibowan Monitoring Stations.
4. The root mean squared standard error of prediction (RMSPE) test is also conducted before and after the intervention. The synthetic control method eliminates the uncertainty of microlevel data in estimating the policy results. A placebo test, as proposed by Abadie et al. (2010), is used to assess the robustness of the COD, NH_3_N, and DO indices. A total of 69 observation points in 2011 that are not involved in the administrative division adjustment are considered in this robustness test. One of the observation points is assumed to undergo an adjustment, and a synthetic version of this region was constructed using other regions outside Hefei. A regression model is applied to the three indicators, COD, NH_3_N, and DO. If the regression results are significant, the difference is not caused by the division adjustment policy but by chance. A large RMSPE indicates that the fitting effect of the synthetic control method may not be appropriate. RMSPE before the intervention is expressed as follows:

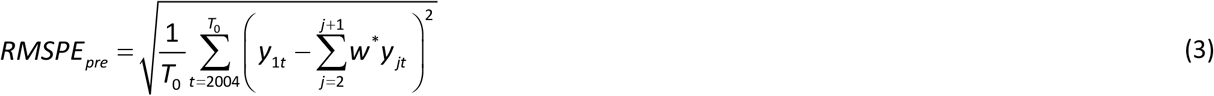 Similarly, the RMSPE after the intervention takes a different time period for the average interval. There are several possible outcomes as follows. If the division adjustment policy has a significant effect in Hefei but not synthetic Hefei, then the model cannot be predicted well and will lead to a larger postintervention RMSPE. As a result, only a placebo effect exists for synthetic Hefei. Moreover, if the results for synthetic Hefei are not well predicted before the intervention, then the preintervention RMSPE will lead to a larger postintervention RMSPE. The ratio of the two can be used to control the impact before the intervention. On the other hand, if the division adjustment has a large policy effect and small placebo effects at other observation points, the ratio of postintervention RMSPE to preintervention RMSPE should be significantly higher in Hefei than at other observation points. Table 8 shows the ratios of RMSPE after the intervention to RMSPE before the intervention for the three indicators at each observation point. As shown in Table 8, for the COD index, the RMSPE values before and after the interventions in Hefei are higher than those of the other 69 observation points, which indicates that the effect of the division adjustment on the COD index is significant. Similar results are observed for the NH_3_N and DO indices. Overall, there is a significant policy effect of the administrative division adjustment of Chaohu, and the synthetic control method has a good fitting effect.
5. A placebo test is performed on each observation point for each indicator. If the observation point has a large RMSPE prior to the division adjustment, then the observation point may not be suitable to construct synthetic Hefei. Therefore, any observation point with an RMSPE greater than 3 times that of Hefei is removed. If values at some observation points are not well synthesized by weighted averages at other observation points, then the difference between the actual and synthetic values may be due to the weighted average. Figure 10 shows the prediction error distribution of COD for Hefei and seven other observation points that meet the requirement. The black line represents the COD prediction error in Hefei, while the gray line represents the COD prediction error for the rest of the observation points. Before 2011, the prediction error of Hefei is moderate. After 2011, there is a significant difference between the true value and the composite value of COD in Hefei city. Only one observation point exceeds Hefei city in 2013 and may be due to the error caused by the weighted average. However, the results show that the division adjustment has a significant negative effect on COD in Hefei.

**Figure 6.**
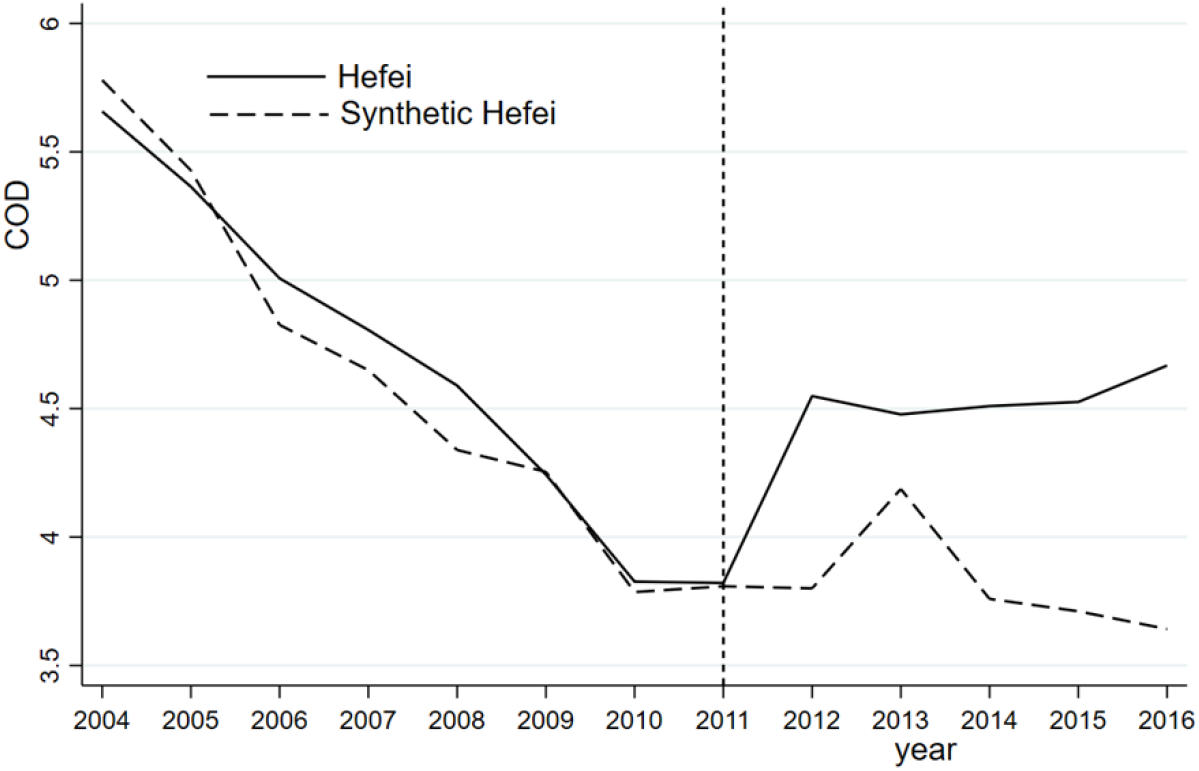
COD of Hefei and synthetic Hefei (excluding Beijing, Shanghai, and Tianjin)

**Figure 7.**
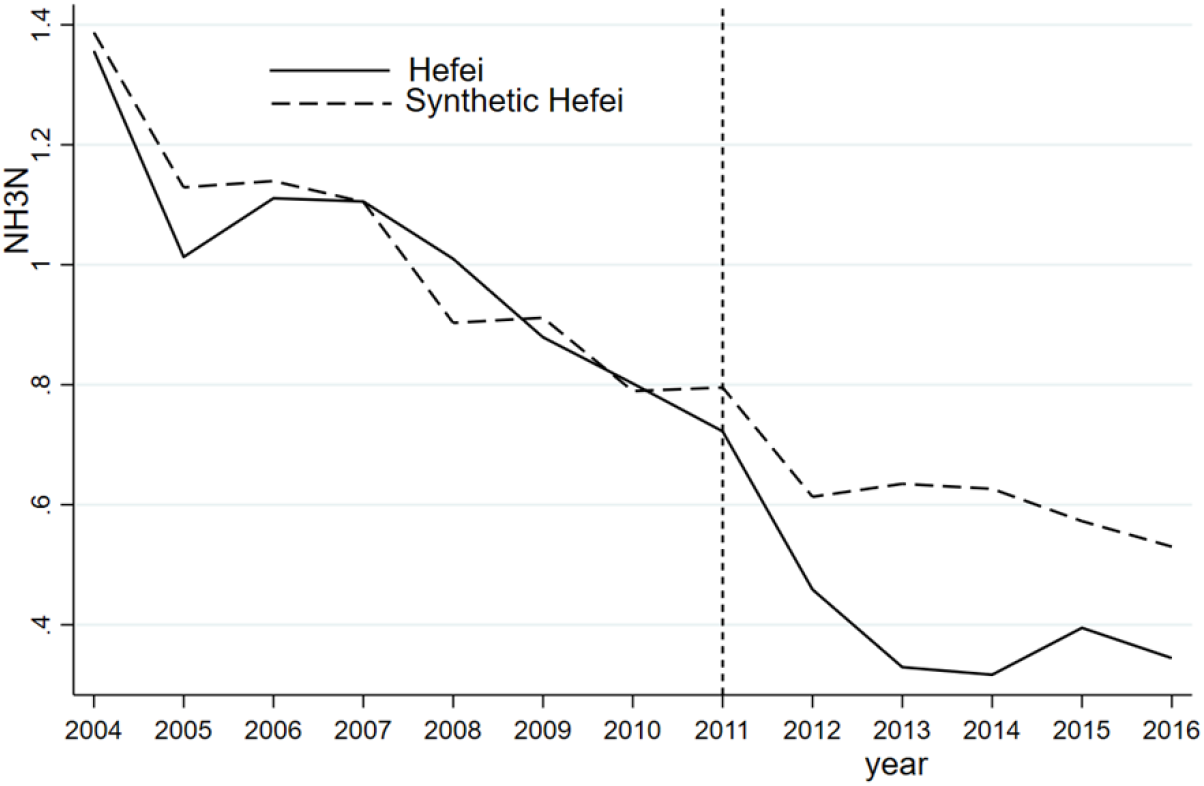
NH_3_N of Hefei and synthetic Hefei (excluding Beijing, Shanghai, and Tianjin)

**Figure 8.**
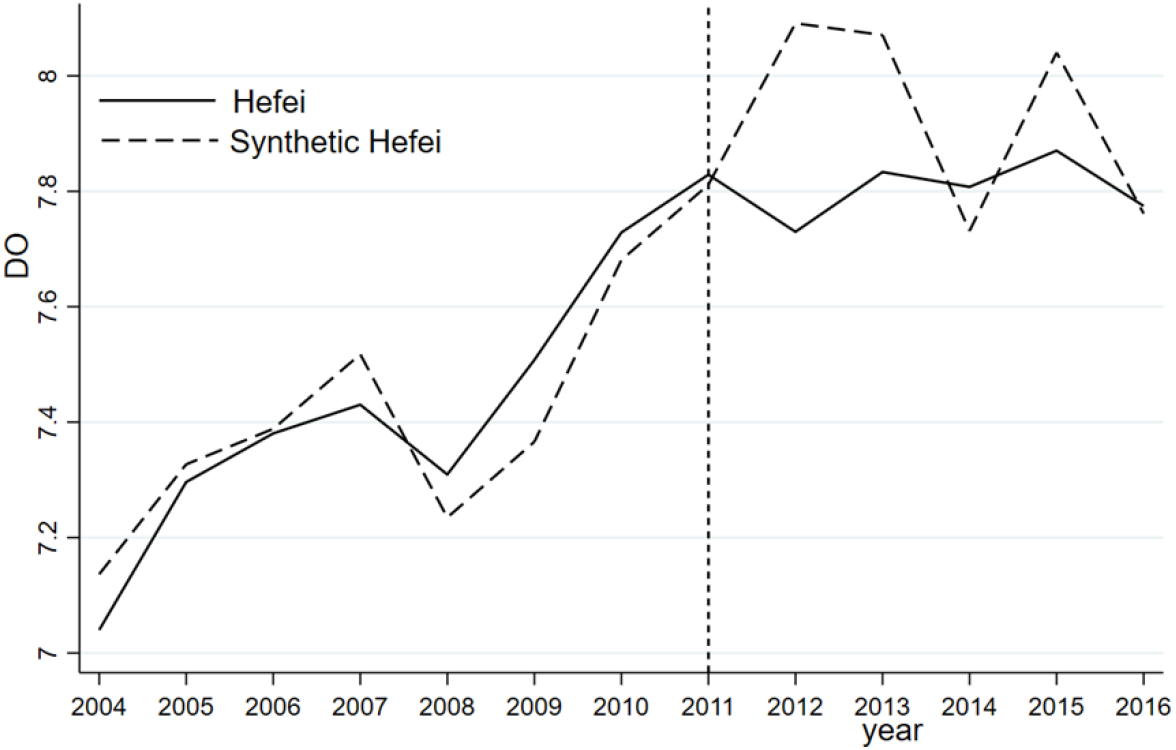
DO of Hefei and synthetic Hefei (excluding Beijing, Shanghai, and Tianjin)

**Figure 9.**
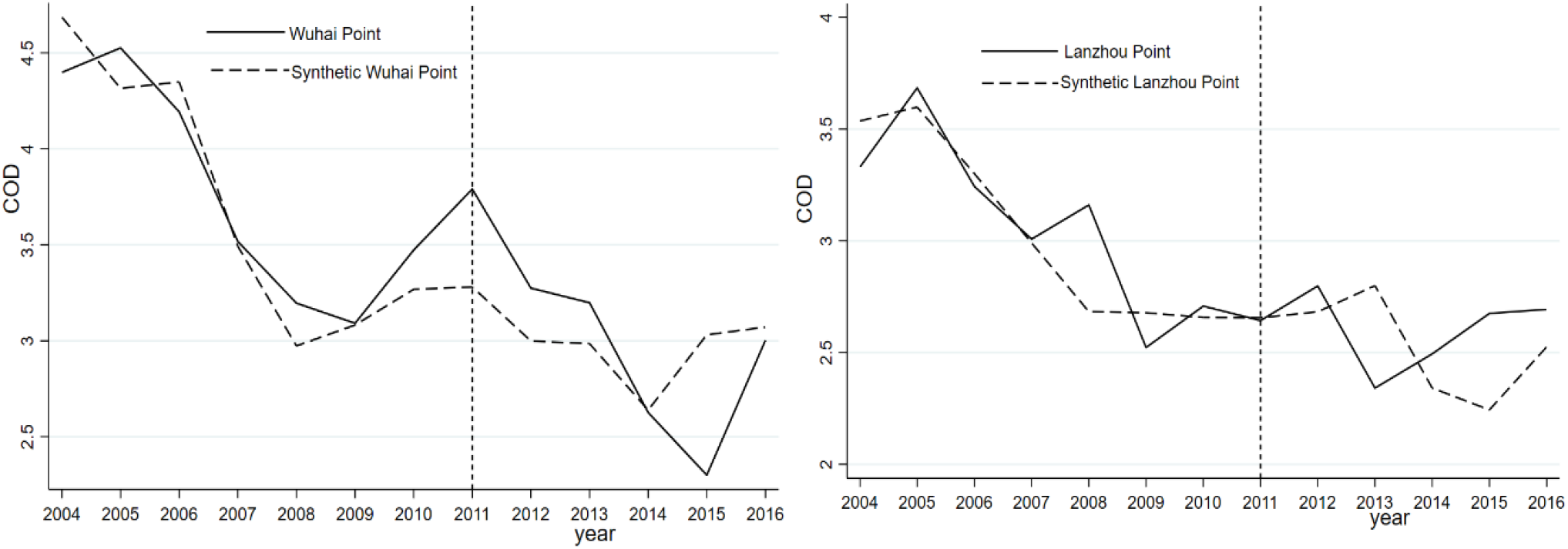
The synthetic effect of COD in the largest and smallest weights in the control group

**Table 8.** RMSPE ratio of the pre- and post-the division adjustment (post-RMSPE /pre-RMSPE)

| ID | Province | City | COD | NH<sub>3</sub>N | DO | ID | Province | City | COD | NH<sub>3</sub>N | DO |
| --- | --- | --- | --- | --- | --- | --- | --- | --- | --- | --- | --- |
| 1 | Shanghai | Shanghai | 0.6734 | 0.3690 | 2.9564 | 29 | Guangdong | Qingyuan | 1.2379 | 0.3010 | 1.1062 |
| 2 | Yunnan | Kunming | 1.4919 | 0.2630 | 0.5278 | 30 | Guangxi | Pingxiang | 0.8242 | 1.3118 | 1.1302 |
| 3 | Yunnan | Kunming | 1.4922 | 0.2630 | 0.5331 | 31 | Guangxi | Nanning | 1.0335 | 0.5667 | 1.4244 |
| 4 | Inner Mongolia | Wuhai | 0.8013 | 0.3600 | 1.8271 | 32 | Guangxi | Guilin | 0.8370 | 0.6049 | 2.2648 |
| 5 | Inner Mongolia | Baotou | 0.5626 | 0.4909 | 2.0934 | 33 | Guangxi | Wuzhou | 0.5575 | 0.2560 | 4.1551 |
| 6 | Beijing | Beijing | 0.7124 | 0.3543 | 1.5660 | 34 | Guangxi | Guigang | 0.8683 | 0.5885 | 0.9247 |
| 7 | Beijing | Beijing | 0.7305 | 0.6873 | 1.9775 | 35 | Jiangsu | Nangjing | 0.5850 | 0.7046 | 1.5708 |
| 8 | Jilin | Baicheng | 2.5117 | 0.5763 | 0.4093 | 36 | Jiangsu | Yixing | 1.8371 | 0.7966 | 1.3845 |
| 9 | Jilin | Changchun | 1.4313 | 0.7768 | 1.6882 | 37 | Jiangsu | Yangzhou | 0.5687 | 0.3569 | 0.4033 |
| 10 | Sichuan | Leshan | 1.0634 | 0.8544 | 1.3291 | 38 | Jiangsu | Wuxi | 1.8371 | 0.7965 | 1.3842 |
| 11 | Sichuan | Yibin | 0.2963 | 0.5953 | 0.4286 | 39 | Jiangsu | Suqian | 0.7012 | 0.3475 | 2.0027 |
| 12 | Sichuan | Panzhihua | 1.0919 | 0.6345 | 0.3912 | 40 | Jiangsu | suzhou | 0.7131 | 0.9520 | 0.7442 |
| 13 | Sichuan | Luzhou | 0.4238 | 0.3856 | 0.8854 | 41 | Jiangsu | Xuzhou | 0.6468 | 0.0219 | 2.2489 |
| 14 | Tianjin | Tianjin | 0.5771 | 1.8599 | 3.7492 | 42 | Jiangxi | Jiujiang | 0.7882 | 0.2799 | 0.4785 |
| 15 | Tianjin | Tianjin | 0.3517 | 0.8844 | 1.6014 | 43 | Jiangxi | Jiujiang | 0.8965 | 0.2419 | 0.4832 |
| 16 | Anhui | Hefei | 3.6169 | 1.7535 | 3.6136 | 44 | Jiangxi | Nanchang | 0.6567 | 0.1337 | 3.3466 |
| 17 | Anhui | Anqing | 0.5591 | 0.4670 | 1.2333 | 45 | Hebei | Zhangjiakou | 0.7341 | 0.1204 | 1.3325 |
| 18 | Anhui | Huaibei | 1.7651 | 0.5411 | 0.5501 | 46 | Hebei | Shijiazhuang | 1.2374 | 0.7338 | 0.8283 |
| 19 | Anhui | Huainan | 1.0810 | 0.2876 | 0.9103 | 47 | Hebei | Zhoukou | 1.1023 | 0.2467 | 0.4273 |
| 20 | Anhui | Fuyang | 0.7814 | 0.4880 | 1.5441 | 48 | Hebei | Zhoukou | 1.1023 | 0.2467 | 0.4273 |
| 21 | Anhui | Bengbu | 0.2824 | 0.5449 | 2.4805 | 49 | Hebei | Jiaozuo | 0.8239 | 0.9146 | 0.8987 |
| 22 | Anhui | Fuyang | 0.7814 | 0.4880 | 1.5441 | 50 | Hebei | Zhumadian | 0.7342 | 0.0923 | 1.3735 |
| 23 | Shandong | Linyi | 2.5305 | 0.1179 | 0.8819 | 51 | Zhejiang | Jiaxing | 0.5423 | 0.8051 | 0.8258 |
| 24 | Shandong | Zaozhuang | 0.3179 | 0.5217 | 1.0110 | 52 | Zhejiang | Hangzhou | 0.4688 | 0.8188 | 0.9832 |
| 25 | Shandong | Jinan | 2.8683 | 1.0434 | 1.0773 | 53 | Zhejiang | Huzhou | 0.1677 | 0.4330 | 0.6828 |
| 26 | Shanxi | Yizhou | 0.2973 | 0.6635 | 0.8535 | 54 | Hubei | Shiyan | 0.6083 | 0.3080 | 0.8944 |
| 27 | Shanxi | Yuncheng | 0.2491 | 0.5905 | 0.6278 | 55 | Hubei | Yichang | 1.2934 | 0.8323 | 0.6172 |
| 28 | Guangdong | Guangzhou | 0.5819 | 0.8724 | 1.2096 | 56 | Hubei | Wuhan | 0.2838 | 0.6456 | 1.2043 |

Table 8 (ctd...). RMSPE ratio of the pre- and post-division adjustment (post-RMSPE /pre-RMSPE)
| ID | Province | City | COD | NH<sub>3</sub>N | DO | ID | Province | City | COD | NH<sub>3</sub>N | DO |
| --- | --- | --- | --- | --- | --- | --- | --- | --- | --- | --- | --- |
| 57 | Hunan | Yueyang | 0.7084 | 0.1993 | 0.9415 | 64 | Liaoning | Yingkou | 0.8573 | 0.6536 | 0.4338 |
| 58 | Hunan | Changsha | 0.6739 | 0.1806 | 0.8526 | 65 | Liaoning | Liaoyang | 0.3909 | 0.4917 | 3.0067 |
| 59 | Hunan | Changsha | 0.6955 | 0.5080 | 1.7045 | 66 | Liaoning | Tieling | 0.2249 | 0.1566 | 0.9563 |
| 60 | Gansu | Lanzhou | 1.4167 | 0.2004 | 1.2052 | 67 | Chongqing | Chongqing | 0.5374 | 0.6881 | 1.2156 |
| 61 | Fujian | Fuzhou | 0.6832 | 0.1025 | 2.7461 | 68 | Heilongjiang | Tongjiang | 0.3880 | 0.9562 | 1.6167 |
| 62 | Liaoning | Fushun | 0.3725 | 0.4147 | 0.3570 | 69 | Heilongjiang | Zhaoyuan | 0.2320 | 0.4491 | 0.8152 |
| 63 | Liaoning | Panjin | 0.1954 | 0.2630 | 1.5290 | 70 | Heilongjiang | Heihe | 1.0721 | 2.1523 | 1.0450 |

**Figure 10.**
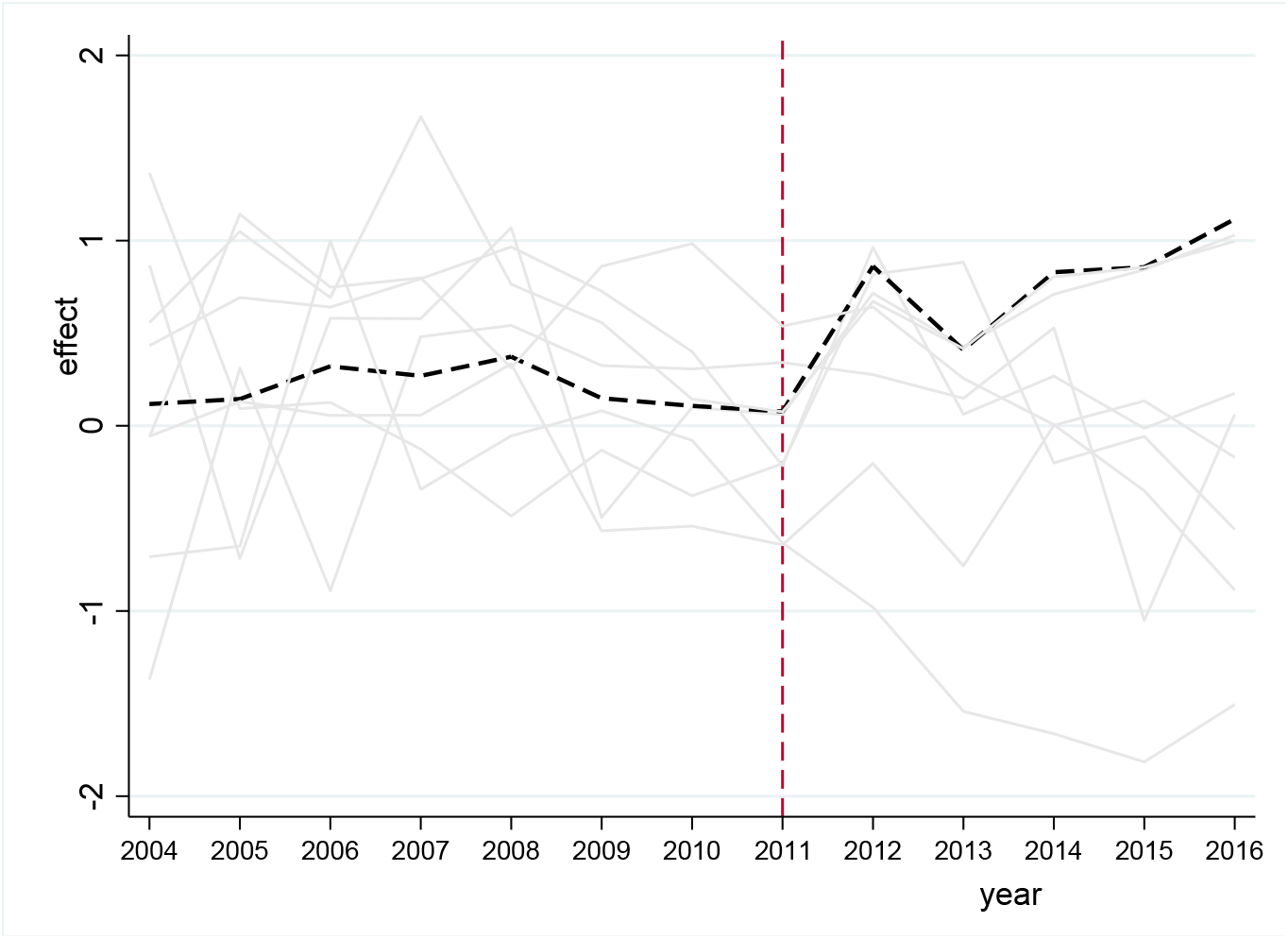
Prediction error distribution of COD for Hefei

Figure 11 shows the prediction error distribution of the NH_3_N index between Hefei and the other 10 observation points. The black line represents the prediction error of COD in Hefei, and the other gray lines represent the error distributions of nine observation points. Overall, the gap between Hefei and the other observation points exhibits a downward trend from 2009 to 2016 and reveals that the administrative division adjustment has an impact on the ammonia nitrogen index of Hefei. In contrast, Figure 12 shows the placebo test results of the DO index in Hefei and 27 other observation points. The black dotted line represents the treatment effect of Hefei (the difference between the DO index of Hefei and that of synthetic Hefei), and the gray solid line represents the placebo effect of 27 observation points (i.e., the difference between the DOs of these observations and their corresponding synthetic observations). Clearly, compared with the placebo effect of other observation points, the DO (negative) treatment effect of Hefei is larger, and the division adjustment of Chaohu has a significant negative effect on the DO index of Hefei.

**Figure 11.**
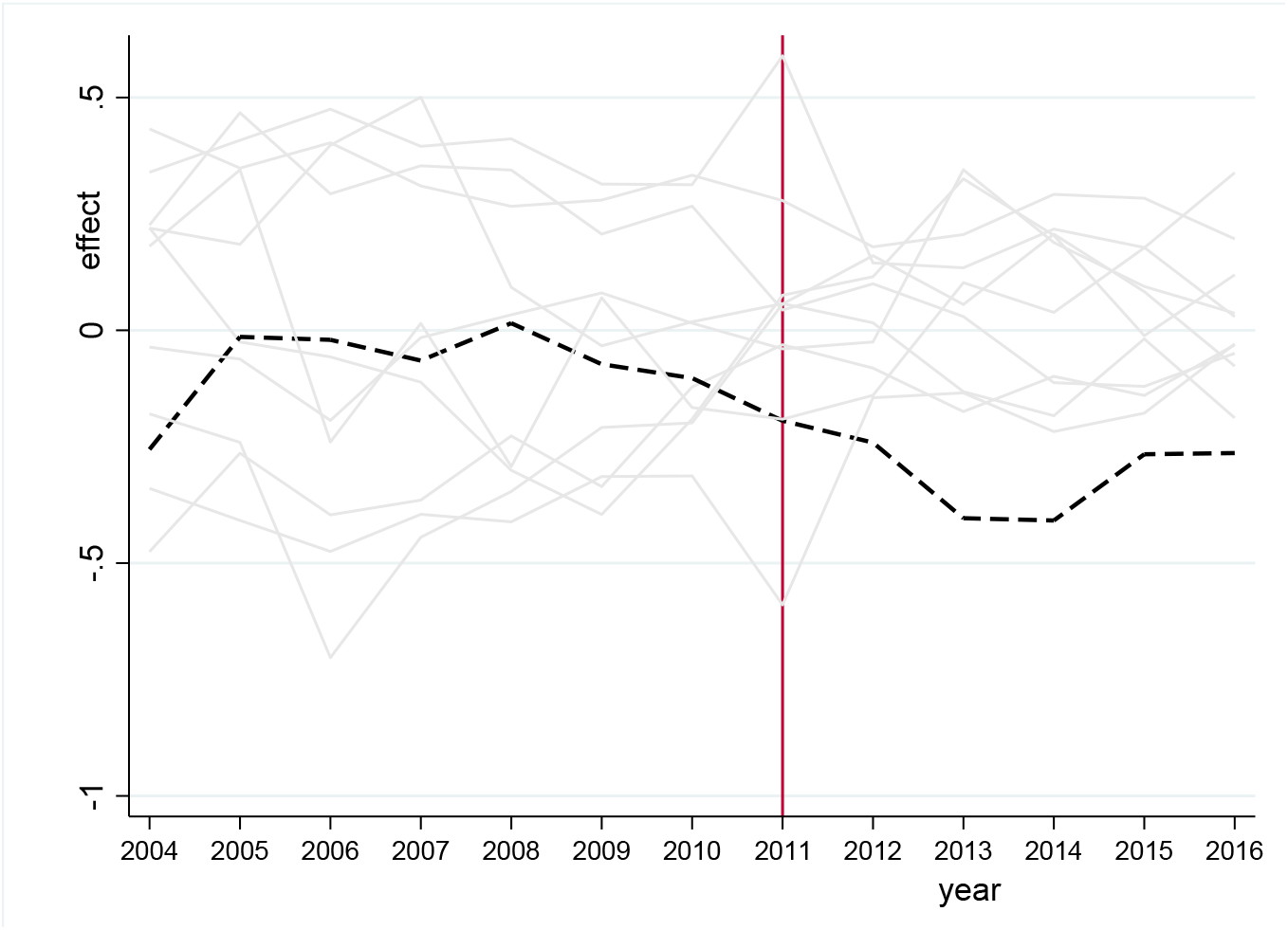
Prediction error distribution of NH_3_N for Hefei

**Figure 12.**
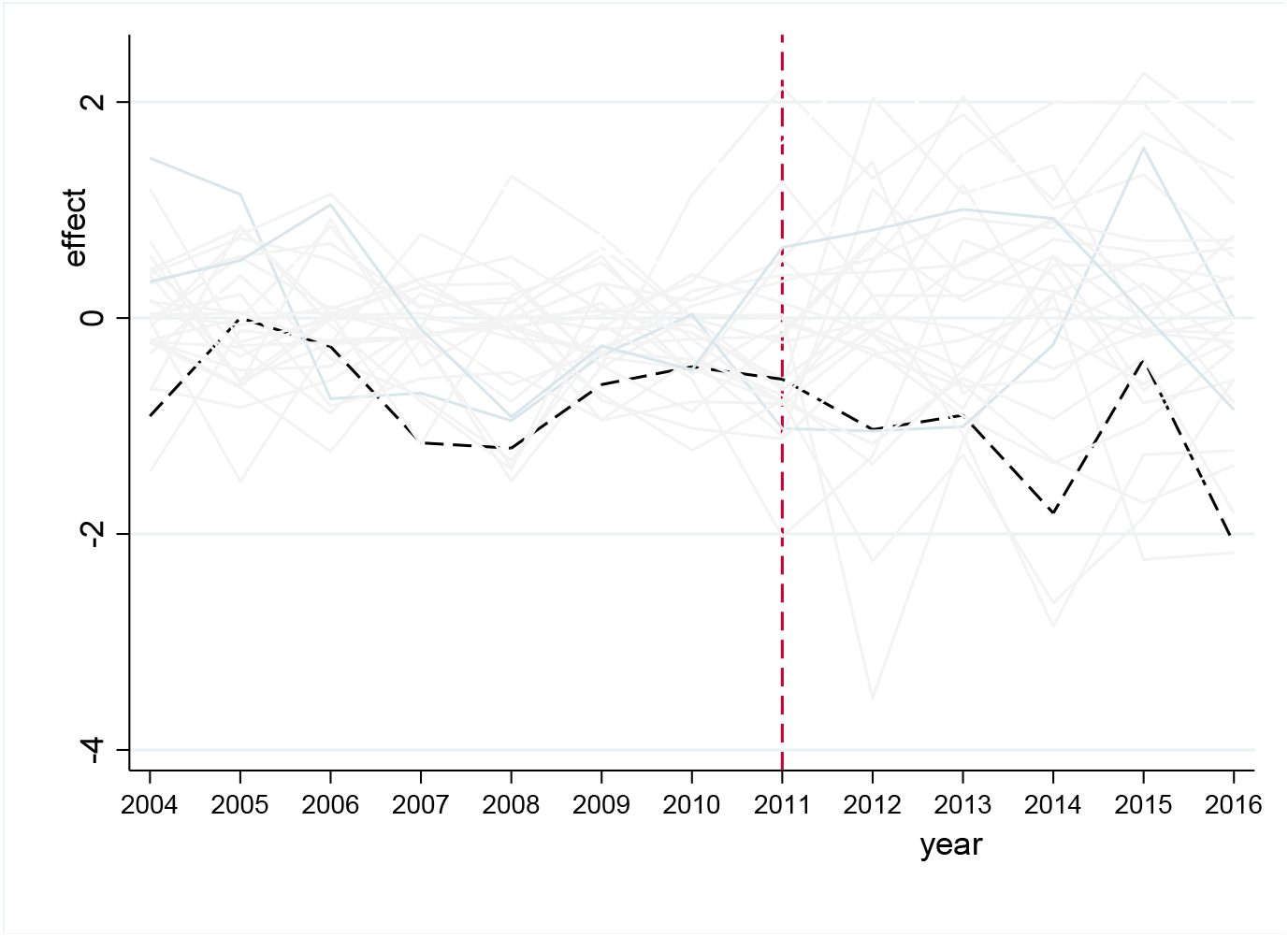
Prediction error distribution of DO for Hefei

### 5.3 Economic and environment effects of administrative division adjustments in Chaohu

The economic and environmental impacts of administrative division adjustments are further analyzed to explain the results.

#### (a) Impact of the administrative division adjustment of Chaohu on Hefei’s economy

A total of 284 prefecture-level cities are used for the synthesis. Table 9 shows that the coefficients of the variables in the synthesis group are similar to those in the experimental group, indicating that the synthesis 22 effect is satisfactory.

**Table 9.** Comparison of treated and synthetic predictors of Log (GDP)

| Predictor Variables | Treated | Synthetic |
| --- | --- | --- |
| LnI | 6.0493 | 5.9450 |
| Lnfix | 7.0365 | 7.0339 |
| Second | 47.3807 | 47.3840 |
| Third | 44.7195 | 44.7208 |
| Labpercent | 62.4605 | 56.4652 |
| Lnpy | 9.9563 | 9.9572 |
| Lnarea | 9.3474 | 9.3470 |
| Lnpop | 6.5209 | 6.5210 |

In table 9, lny is the natural logarithm output of synthetic Hefei, lnl is the labor force, lnfix is a fixed asset investment, second denotes the proportion of secondary industry, third denotes the proportion of the tertiary industry, labpercent is the employment level, lnpy is the GDP per capita, lnarea is the city area, and lnpop is the urban population. The sample data are processed to ensure that the variables used for synthetic control are close to those of Hefei before the division adjustment.

The results indicate that GDP of synthetic Hefei will continue to grow after 2011. Figure 13 shows that Hefei’s GDP experiences a short-term downward trend in 2011 because the prefecture-level municipality of Chaohu is dominated by agriculture, has slow development and a large population, and its overall economic level lags far from that of Hefei city. After the abolition of the prefecture-level municipality of Chaohu, Lujiang county and merger with Chaohu city, the economic system of Hefei impacts the economy of Hefei. However, this change was not continuous, and there was a temporary decline in the output of Hefei.

**Figure 13.**
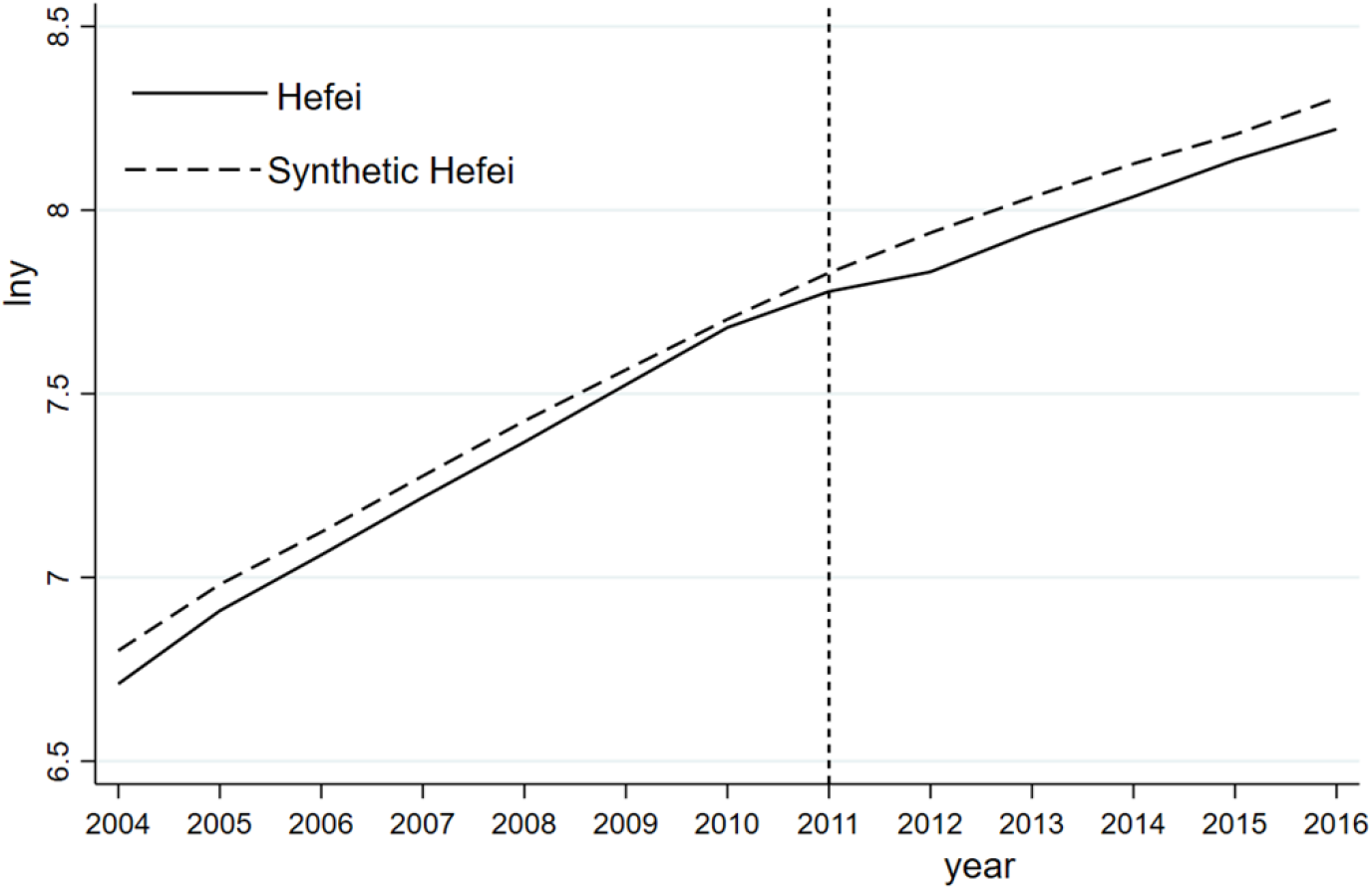
Log (GDP) of Hefei and synthetic Hefei

To clearly illustrate the impact of the abolition of the prefecture-level municipality of Chaohu, a synthetic forecast of the GDP of the secondary industry is conducted. The synthetic results are shown in Table 10. Except for the proportion of the tertiary industry, the synthetic errors of the indicators indicate that the fitting effect is good and accurately simulates Hefei without division adjustment. In contrast, as shown in Figure 14, the trend of the GDP in the secondary industry contradicts the overall economic trend. Industrial development of Hefei has surpassed that of synthetic Hefei, which indicates that the division adjustment has a significant driving effect on the industrial development of Hefei. Overall, administrative division adjustment has slowed the overall economic development rate of Hefei but improved industrial growth, which is consistent with Hypothesis II. Regional growth and Chaohu governance are more focused on economic growth, especially industrial expansion, weakening Chaohu governance.

**Table 10.** Comparison of treated and synthetic predictors of Log (GDP industry)

| Predictor Variables | Treated | Synthetic |
| --- | --- | --- |
| *LnI* | 6.0493 | 6.0368 |
| *Lnfix* | 7.0365 | 7.0101 |
| *Second* | 47.3807 | 47.3822 |
| *Third* | 44.7195 | 41.9444 |
| *Labpercent* | 62.4605 | 62.4595 |
| *Lnpy* | 9.9563 | 9.9564 |
| *Lnarea* | 9.3474 | 9.3469 |
| *Lnpop* | 6.5209 | 6.5220 |

**Figure 14.**
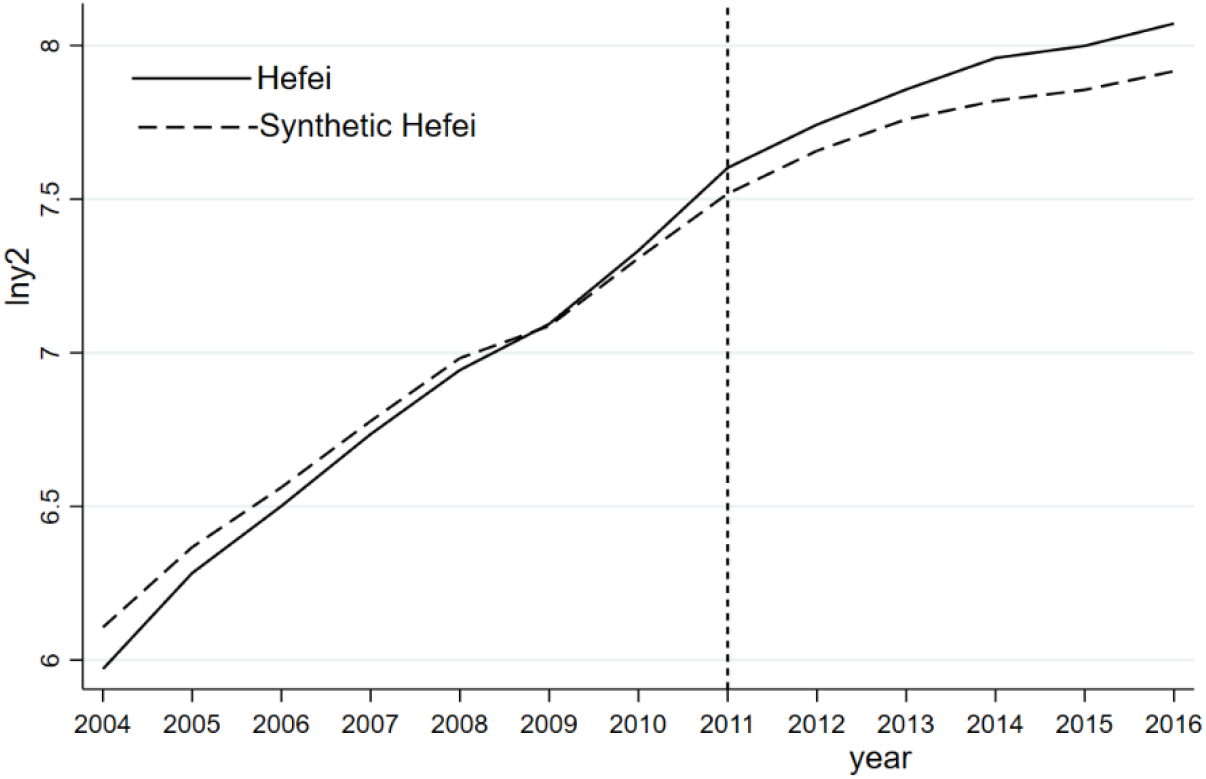
Log (GDP industry) of Hefei and synthetic Hefei

#### (b) NH_3_N, COD, and DO indices

The decline in NH_3_N may be mainly due to industrial development replacing agricultural development. A series of industrial expansions, such as urban construction and real estate development, have caused a reduction in agricultural production. The chemical fertilizer and pesticide use in Hefei show a significant decline after the division adjustment in 2011. Although the amount of agricultural fertilizer used by Hefei has shown a downward trend since 2006, the decline is even greater in 2011, reaching 40.12%. Figure 15 indicates that the amount of pesticide used is near 300 tons in 2006-2010 and then undergoes a large decline, reaching 18.23% in 2011, with a continued downward trend after 2011. The reduction in the scale of agriculture leads to a decline in the use of chemical fertilizers and pesticides. Therefore, development of agricultural production is crowded out by industrial development, which has led to a decline in the NH_3_N index.

**Figure 15.**
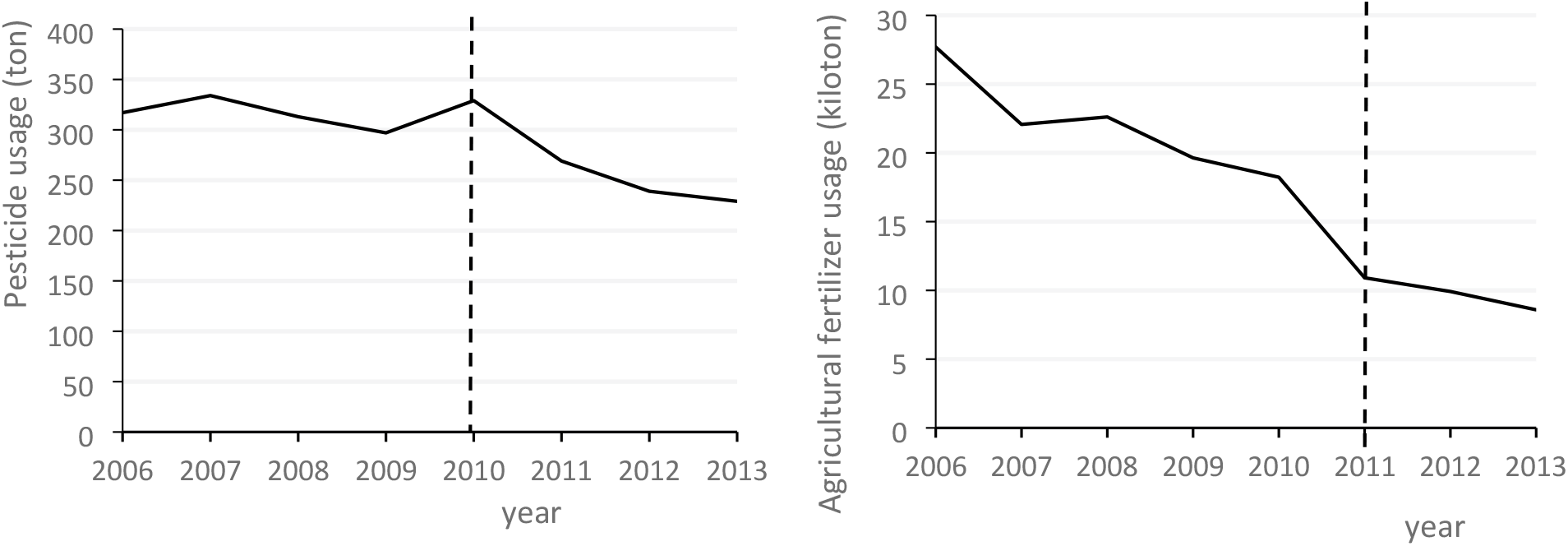
Pesticide and fertilizer consumptions in Hefei

Deterioration of the COD and DO is related to the rapid development of industry and urbanization. A large amount of domestic sewage and industrial sewage are discharged without effective treatment, resulting in an increase in the amount of water pollution in Chaohu Lake. Many sewage treatment plants have not been completed on time, leading to direct discharge of domestic sewage. For instance, low sewage treatment rate and lack of treatment facilities have led to poor results. The industrial water intensity has been the main obstacle to the ecological security of the eastern half of Chaohu Lake in recent years, while the pollution of the western half of the lake mainly comes from urbanization development. The wetland area, domestic water intensity, and artificial forestation area are the main factors affecting the ecology of Chaohu Lake (Tang et al., 2019). Deterioration of the COD and DO indicators after the division adjustment is closely related to the intensive urbanization and industrialization development in Hefei.

Several reasons for the poor environmental governance of Chaohu Lake after the division adjustment can be summarized as follows:

a. In terms of development space, urbanization construction has taken over ecological land, and a large number of illegal constructions have occurred. As stated by the central inspection team, as an expanded urban space, Hefei has seized a large amount of ecological land for commercial development, tourism development, and urban construction. Numerous commercial and real estate projects are illegally approved and constructed in the Chaohu Basin water environment protection area. Moreover, several problems have been experienced in Chaohu; for instance, the wave proof forest platform is damaged and aquatic plants are completely destroyed, which will lead to extinct ecological functions. In the name of restoration, tourism development in Chaohu Lake continuously reduces wetlands and supports tourism development. The destruction of ecological wetlands reduces the self-recovery function and recovery of the lake.
b. The poor management of Chaohu Lake is another major problem. A specialized organization for the management of Chaohu Lake is established in 2012 to provide unified protection and supervision. However, the Chaohu Administration Bureau is not performing its environmental protection responsibilities, partly because of the mismatch of financial power and administrative power, for instance, lake invasion and wetland destruction prevention.
c. The emphasis of the local government is more on the economy instead of the environment, resulting in environmental regulations and pollution control projects that are not implemented on schedule. For example, the Anhui Provincial Government adjusts the weights of target management assessments in prefecture-level cities in 2016. The weight of economic development rises from 14.6% to 22.3% in the previous year to 27.5% to 32.5%. However, the evaluation weight of ecological and environmental indicators changes from 14.6%-22.3% to 13.5%-20.5%. The deviation has led to economic development but poor environmental protection. Some government agencies are more inclined to engage in economic development and ignore environmental protection issues. For instance, the requirements stipulated in the “Chaohu Basin Water Pollution Prevention Regulations” have not been implemented, and the third phase of the Shiwili River Sewage Treatment Plant, initiated in 2013, has not been completed, resulting in approximately 60,000 tons of sewage being discharged into Chaohu Lake. Water pollution treatment in Chaohu Lake, which will affect the sustainable development of the economy, increase the pressure on the environment, and lead to the improvement of water quality, is usually placed after economic development in terms of importance.
d. A limited role of nonprofit organizations and public participation are used to explain the governance of Chaohu Lake. Learning from the experience of large lakes in Western countries, active public participation is required to obtain good lake water pollution treatment results. The public participates in pollution control in a variety of ways, such as by participating in the drafting of water pollution control policies and building their own awareness of ecological environment protection. Moreover, Lake Biwa in Japan is another good example, where nonprofit organizations and the public are playing important roles in the prevention and control of water pollution. Although the government always plays a dominant role in the treatment of lake pollution, the government alone is insufficient to overcome the water pollution problem. A collaboration between enterprises, nonprofit organizations, the public, and governance entities is required to establish a reasonable governance cooperation mechanism for managing Chaohu Lake.

## 7. Conclusions

Administrative division adjustment has various social, economic, and environmental effects. Previous literature has focused on the economic effects arising from administrative division adjustment and has ignored the environmental effects. The division adjustment in Chaohu is conducive to environmental governance in the long term. The synthetic control method is used to identify the effect of the administrative division adjustment on the water quality of Chaohu Lake. After the division adjustment, some water quality indicators such as NH_3_N have indeed been alleviated and dropped by approximately 30%; however, other major pollution indicators, such as COD and DO, have deteriorated to varying degrees. For instance, COD has increased by approximately 25%. After excluding municipalities from the sample, we compare the mean square prediction error before and after the division adjustment. A placebo test applied to assess the robustness of the results indicates that the environmental policy effect of the division adjustment is not satisfactory. Furthermore, several factors have influenced economic and industrial expansion, including the rapid expansion of urban areas, the imperfect management system of Chaohu Lake, and the emphasis on the economic performance of officials.

Several important policy implications can be drawn from the current study. First, returning to the initial intention of administrative division adjustment, the main reason the prefecture-level municipality of Chaohu is split and merged is to improve the governance of Chaohu Lake. Prior to 2011, Chaohu Lake has the most unsatisfactory governance of the five largest freshwater lakes in China. The central government is determined to improve the management system of Chaohu Lake by dismantling and merging Chaohu city to achieve better environmental governance and economic development. Therefore, we re-emphasize and rationalize the governance mechanism to achieve the policy objectives of the division adjustment plan. Second, we shift the focus to high-quality development instead of economic development. To achieve this goal, we must not habitually rely on economic growth and environmental governance. In addition, it is necessary to clarify the power and responsibility for environmental monitoring in the Chaohu Lake Basin. For instance, the environmental monitoring and enforcement functions should be the responsibility of the Chaohu Administration Bureau.

## Footnotes

① Information source: http://www.gov.cn/hudong/2017-07/30/content_5214708.html gives way to economic development in Chaohu.

2 The Ninth Five-Year Plan projects are mostly small-scale projects. After the Tenth Five-Year Plan period, only projects with an investment of more than 10 million yuan are counted.

